# Functional Profiling of a Human Odorant Receptor Using Olfactory Cilia Links its Activation to Odor Quality

**DOI:** 10.64898/2026.09.23.753647

**Authors:** Eugene Lempert, Masayo Omura, Raena Mina, Paul Feinstein

## Abstract

The mechanism governing singular odorant receptor (OR) gene choice restricts each olfactory sensory neuron (OSN) to expressing a single OR allele from a repertoire of more than 1,100 receptor genes. Consequently, only a small fraction of OSNs express any given OR, limiting studies of OR function, axonal wiring, and odor coding. Here, we show that multimerizing a 21-bp homeodomain enhancer dramatically increases the probability of receptor choice. We engineered a single-copy Olfr151 minigene carrying a 9×21 enhancer and the human odorant receptor OR10G4, integrated at a genomic site distant from all endogenous OR clusters. This transgene drives singular OR10G4 expression in approximately 75% of OSNs and is accompanied by widespread reductions in endogenous Class I, Class II, and TAAR receptor transcripts and proteins, indicating that cis-regulatory elements govern the probability of OR selection. The abundance of OR10G4-expressing cilia enabled highly sensitive ligand screening using CELIA (Cilia-based Evaluation of Ligands and receptor InterActions). Profiling guaiacol derivatives revealed a strong correspondence among receptor activation, cAMP production, and published human psychophysical responses. Although OR10G4 responds to guaiacol-related odorants, multiple canonical vanilla odorants failed to activate the receptor, arguing against a primary role in vanilla perception. Instead, the lowest EC50 ligands were enriched for compounds human subjects described as smoky. Together, these findings link activation of a single human odorant receptor to a defined perceptual odor-quality dimension and establish a general strategy for decoding the molecular basis of human odor perception.

## Introduction

The olfactory system provides a remarkable model for studying selective molecular interactions. In mature olfactory sensory neurons (mOSNs), a single odorant receptor (OR) allele is actively transcribed from a repertoire of more than 2,000 evolutionarily related alleles. Axons of neurons expressing the same OR preferentially interact and coalesce into homogeneous glomeruli within the olfactory bulb(1–3). At the molecular level, each OR is activated by a distinct subset of ligands that share combinations of physical, chemical, and signaling properties. Although these principles are well established, the OR gene family and its unique cellular host, the OSN, provide an unparalleled experimental system for investigating the mechanisms that govern gene choice, neuronal wiring, and sensory coding.

The singular OR expression, in which each OSN expresses only one OR allele, is fundamental to olfactory function. Despite decades of investigation, several major questions remain unresolved. How does an OSN select a single OR allele from a repertoire of more than 2,000 alleles? How do neurons expressing the same receptor recognize one another and coalesce their axons into receptor-specific glomeruli? And what are the ligand repertoires and perceptual functions of the approximately 400 human OR genes? Resolving these questions is essential for understanding how odor perception emerges from receptor activation. Progress has been limited because a typical OR is expressed in only ∼5,000 OSNs in mice(4–6), making mechanistic studies of individual receptors and receptor-dependent processes inherently difficult(1, 2). We reasoned that addressing all three questions require the ability to drive expression of selected ORs in substantially larger populations of OSNs.

Through a decades-long dissection of olfactory promoter architecture, we identified two critical transcription factor binding motifs involved in OR gene choice: one recognized by Olf1/Ebf1 and a second recognized by the LIM-homeodomain transcription factor LHX2(7–9). Analysis of more than 30 OR minigenes led to the discovery of a potent 21-bp homeodomain enhancer that dramatically increases OR choice frequency when multimerized four or five times (4×21 or 5×21) and placed upstream of an OR transgene (4, 10–14). We subsequently showed that tandem arrays of 5×21-enhanced OR transgenes remain robustly expressed even when integrated far from any of the 30 endogenous OR-containing loci and the eight isolated OR genes that reside outside OR clusters, suggesting that these transgenes function independently of local OR super-enhancer environments(4). Whereas endogenous OR alleles are typically expressed in several thousand neurons, 4×21- and 5×21-enhanced Olfr151 transgenes can be expressed in populations one to two orders of magnitude larger while preserving singular receptor expression(4). Initially, we quantified these effects by extrapolating from glomerular size to OSN number. Increasing the number of neurons expressing a single OR creates opportunities to investigate OR gene choice, identify OR-dependent growth cone interactions that mediate homotypic axonal sorting, and generate abundant populations of olfactory cilia expressing a common receptor.

To further increase receptor choice frequency, we generated OR transgenes carrying nine tandem copies of the 21-bp enhancer (9×21). We identified a 9×21-OR10G4 transgenic mouse line carrying a single-copy integration on chromosome 3 more than 32 Mb from the nearest endogenous OR cluster. In this line, OR10G4 is expressed in approximately 75% of the cells of olfactory sensory neuron lineages in the olfactory epithelium, and this extensive expression is accompanied by reductions in endogenous Class I OR, Class II OR, and TAAR transcripts, suggesting that the transgene participates in or competes for a common OR choice mechanism across all receptor-expressing OSNs. To our knowledge, the only previously reported method capable of producing similarly widespread OR expression is the tTA/TetO system, whose mechanism remains unclear(15–17).

Current models of singular OR choice propose that receptor selection is driven by competition among multi-enhancer assemblies termed Greek Island (GI) hubs, each composed of aggregates of 5–6 enhancers selected from more than 180 Greek Islands distributed throughout OR clusters across nearly all chromosomes(18). Greek Islands contain paired homeodomain (HD) and Olf1/Ebf1 (O/E) motifs and are thought to function critically in cis while acting redundantly in trans to support expression of a chosen OR allele(18–20). Consistent with this interpretation, deletion of individual Greek Islands preferentially affects nearby OR genes while more distal ORs within the same cluster remain largely unaffected(18). In contrast, the robust expression of our single-copy OR10G4-transgene, aided by a compact 189-bp HD-only enhancer and located far from any endogenous Greek Island, provides compelling evidence that an OR locus can effectively nucleate its own choice. These findings suggest that the strength of a choice-promoting enhancer strongly determines the probability that an OR gene will be selected for expression(10).

We previously used ×21-enhanced OR transgenes and isolated olfactory cilia preparations to generate comprehensive ligand profiles for the human odorant receptors OR1A1 and OR5AN1(11). To extend this approach with our Cilia-based Evaluation of Ligands and receptor InterActions (CELIA)(11), we selected OR10G4 because it responds to guaiacol(21), a key constituent of smoky aromas, and because human psychophysical studies have extensively characterized the perceptual properties of guaiacol derivatives(22, 23). Using the OR10G4 transgenic line, we demonstrate that perceptual detection thresholds for halogenated or alkylated derivatives of guaiacol correlate closely with receptor activation measured in native olfactory cilia. These findings support assigning a smoky perceptual quality to OR10G4 and provide a direct link between activation of a single human odorant receptor and a defined odor-quality dimension.

## Results

### OR10G4-expressing transgenic lines

We previously developed a transgenic strategy to increase the number of olfactory sensory neurons (OSNs) expressing a selected odorant receptor (OR). Briefly, we identified a core homeodomain sequence within the OR enhancer H, multimerized it nine times as a 21-bp element (9×21), and placed it upstream of an Olfr151 transgene cassette containing an OR-IRES-reporter construct(11). Whereas an endogenous OR is typically expressed in approximately 5,000 OSNs, this approach increases the number of neurons selecting the transgenic OR by 8- to 100-fold(4, 11).

Using this strategy, we generated two independent transgenic lines carrying the human OR construct *OR10G4-IRES-MylParmGCaMP6f,* which was confirmed by genotyping screening (**Figures 1A, S1A and S1B**). Initial characterization of reporter fluorescence revealed an unexpected difference between the lines. Line 10 (OR10G4#10) exhibited robust fluorescence throughout the olfactory epithelium, whereas Line 7 (OR10G4#7) displayed little to no detectable reporter signal (**Figure S1D**). Because we previously demonstrated that isolated olfactory cilia contain all the molecular machinery required for odorant-induced cAMP production through the canonical OR/G-protein/Adcy3 signaling cascade(11), we tested cilia isolated from both lines using a cAMP-based receptor activation assay as in Omura et al., 2022. Unexpectedly, the canonical OR10G4 agonist guaiacol elicited a substantially stronger response in OR10G4#7 than in OR10G4#10 (**Figure S1C**).

**Figure 1:**
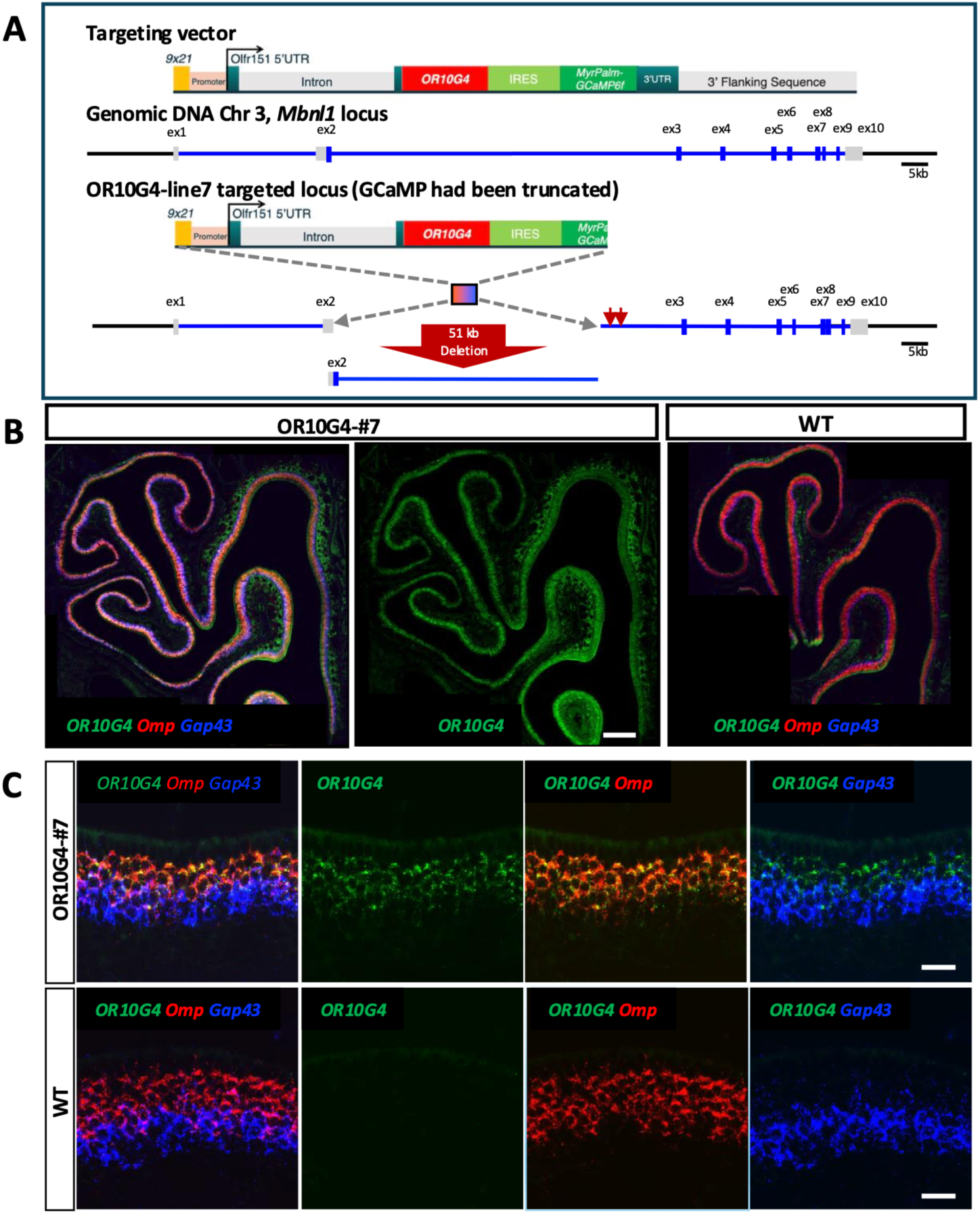
OR10G4 is broadly expressed in Olfactory Sensory Neurons. (**A**). Human OR10G4-expressing transgenic mouse lines were generated by pronuclear injection of a targeting vector containing *9×21* (9 multimers of 21mer high-probability choice element) and the human OR10G4 coding sequence, followed by *IRES-MylPamGCaMP6f*, with the *Olfr151* 5 ‘promoter region and 3 ‘untranslated region (UTR) flanking the backbone. In the highly expressing line #7, the OR10G4 targeting vector was integrated into the *Mbnl1* locus on chromosome 3, with a 51 kb deletion that included part of exon 2 and an intron of *Mbnl1*. Targeted locus amplification (TLA by Cergentis) analysis also showed that the targeting vector was truncated in the middle of the *GCaMP6f* gene, and the mouse line does not have functional GCaMP6f expression (no GCaMP fluorescence) nor *Olfr151-3’UTR*. (**B**). RNAscope on the coronal sections of the olfactory epithelium with *OR10G4*, *Omp*, and *Gap43* probes shows high and broad expression of OR10G4 in the transgenic line, while there is no *OR10G4* expression in the wild-type control. Scale bar 200 µm. (**C**). High-magnification images of RNAscope with *OR10G4*, *Omp*, and *Gap43* probes. *OR10G4* expressed in *Omp-* and *Gap43*-positive cells. Scale bar 20 µm.

To understand this discrepancy, we mapped the transgene integrations by PCR and identified a truncation of the GCaMP portion of the construct in OR10G4#7 (**Figure S1A-1B**). We then performed RNAscope using probes against both the OR10G4 and GCaMP coding sequences. Consistent with the PCR results, OR10G4#7 lacked signal corresponding to the 3′ region of the GCaMP transcript. Surprisingly, however, OR10G4 mRNA was more broadly distributed in OR10G4#7 than in OR10G4#10, although the signal intensity per cell appeared lower (**Figure S1E**).

PCR analysis further suggested that OR10G4#7 contains a single integration of the truncated transgene. Because the integration showed properties consistent with a potential safe-harbor locus, we submitted the line to The Jackson Laboratory for colony preservation. We then compared the original colony (Line 7) with the JAX-derived colony (Line 7J) using RT-qPCR. Assays targeting the OR10G4 coding sequence and the retained 5′ portion of GCaMP confirmed that Line 7J faithfully reproduced the original Line 7 genotype and expression profile. Both Line 7 and Line 7J contained substantially higher levels of transgenic mRNA than Line 10 (**Figure S1F**).

To determine whether the transgene co-opted the endogenous mechanism of singular OR gene choice rather than being expressed alongside native receptors, we quantified expression of seven endogenous OR genes by RT-qPCR. OR10G4#7 and OR10G4#7J exerted a profound suppressive effect on endogenous OR expression, reducing transcript abundance to less than 25% of wild-type levels for six of seven tested receptors. In contrast, OR10G4#10 reduced expression of only one of seven ORs to a comparable extent (**Figure S1G**). RNAscope further showed that OR10G4 expression in Line 7 was nearly ubiquitous, with near-complete overlap between OR10G4 mRNA and the mature OSN marker Omp (**Figure 1B-1C**). Together, these findings indicate that the OR10G4#7 transgene effectively commandeers the endogenous mechanism of singular OR gene choice, driving OR10G4 expression in most OSNs while simultaneously suppressing expression of the native OR repertoire. As a result, OR10G4#7 produces an almost monoclonal olfactory epithelium and provides a powerful model for investigating OR choice, receptor function, and odor coding.

### Identification of the OR10G4#7 Integration Site

Establishing a genomic locus that reliably supports high-frequency expression of ×21-enhanced OR transgenes would greatly facilitate generating a library of mouse lines in which a chosen OR coding sequence predominates throughout the olfactory epithelium (**Figure 7C**). To identify the integration site responsible for the robust OR10G4#7 phenotype, we used a strategy called Targeted Locus Amplification (TLA) sequencing. TLA analysis revealed that OR10G4#7 contains a single-copy transgene integration on chromosome 3, with no evidence of transgene concatemerization or end-to-end joining events. The transgene is inserted within the first exon of a presumptive third transcriptional start site of the *Mbnl1* gene and is associated with an approximately 50-kb deletion encompassing both exonic and intronic *Mbnl1* sequences (**Figure 1A**; Cergentis Report of TLA data available upon request).

The absence of GCaMP fluorescence and the failure to detect full-length *GCaMP* transcripts by RNAscope are explained by a deletion at the 3′ end of the integration cassette. This deletion originates within the *GCaMP* coding sequence and extends through the entire 3.5-kb Olfr151 3′ UTR and polyadenylation region, fully consistent with our PCR-based mapping results. Despite this truncation, the OR10G4 coding sequence remains intact and is expressed at levels sufficient to drive the dominant OR10G4 phenotype.

Notably, chromosome 3 contains only two functional OR genes (*Olfr1402* and *Olfr266,* separated by 8Mb) and their associated Greek Island enhancers (**Folegandros** and **Kos**, respectively), all located more than 30 Mb from the *Mbnl1* locus(19). Consequently, the OR10G4#7 integration resides outside any genomic region previously associated with OR super-enhancer activity or OR-specific regulatory architecture. The ability of a single-copy 9×21-enhanced transgene to dominate OR choice from this location demonstrates that neither proximity to an OR cluster nor association with a known Greek Island is required for robust receptor selection, making it an attractive candidate site for future engineering of receptor-specific mouse lines.

### OR10G4 Expression Suppresses the Endogenous OR Repertoire

The widespread expression of *OR10G4* and the robust guaiacol-induced responses observed in the OR10G4#7 line, together with the reduction in endogenous OR transcripts detected by RT-qPCR, prompted us to investigate the transgene’s impact on global OR gene expression. Bulk RNA-seq of the olfactory epithelium revealed a dramatic effect: nearly every endogenous OR gene showed significantly reduced expression, defined as p ≤ 0.05 and at least a one-fold change in transcript abundance (**Figure 2A**). These findings were highly consistent with our targeted RT-qPCR analyses **(Figure S1G, S1H, and Table S1**). As expected, *Olfr151*, which shares the same 5′ regulatory region as the OR10G4#7 transgene, showed elevated expression (**Figure 2A and 3A**). In contrast, expression of *Ms4a* and *Gucy* family genes, which mark non-OR sensory cell populations in the olfactory epithelium, were unaffected (**Figure 3B**). Notably, Class I ORs, Class II ORs, and TAAR genes, which define three distinct classes of chemosensory neurons, all exhibited substantial reductions in expression with no obvious chromosome-specific exceptions (**Figure 2B**). Although suppression was observed across all chromosomes, analysis by dorsal-ventral index (DVI, (4)) revealed a significant spatial bias: *OR* genes expressed in ventral zones were more strongly suppressed than those expressed dorsally, and the distributions of ventral and dorsal *OR* gene log2 fold-changes differed significantly (**Figures 2C and 3C).**

**Figure 2:**
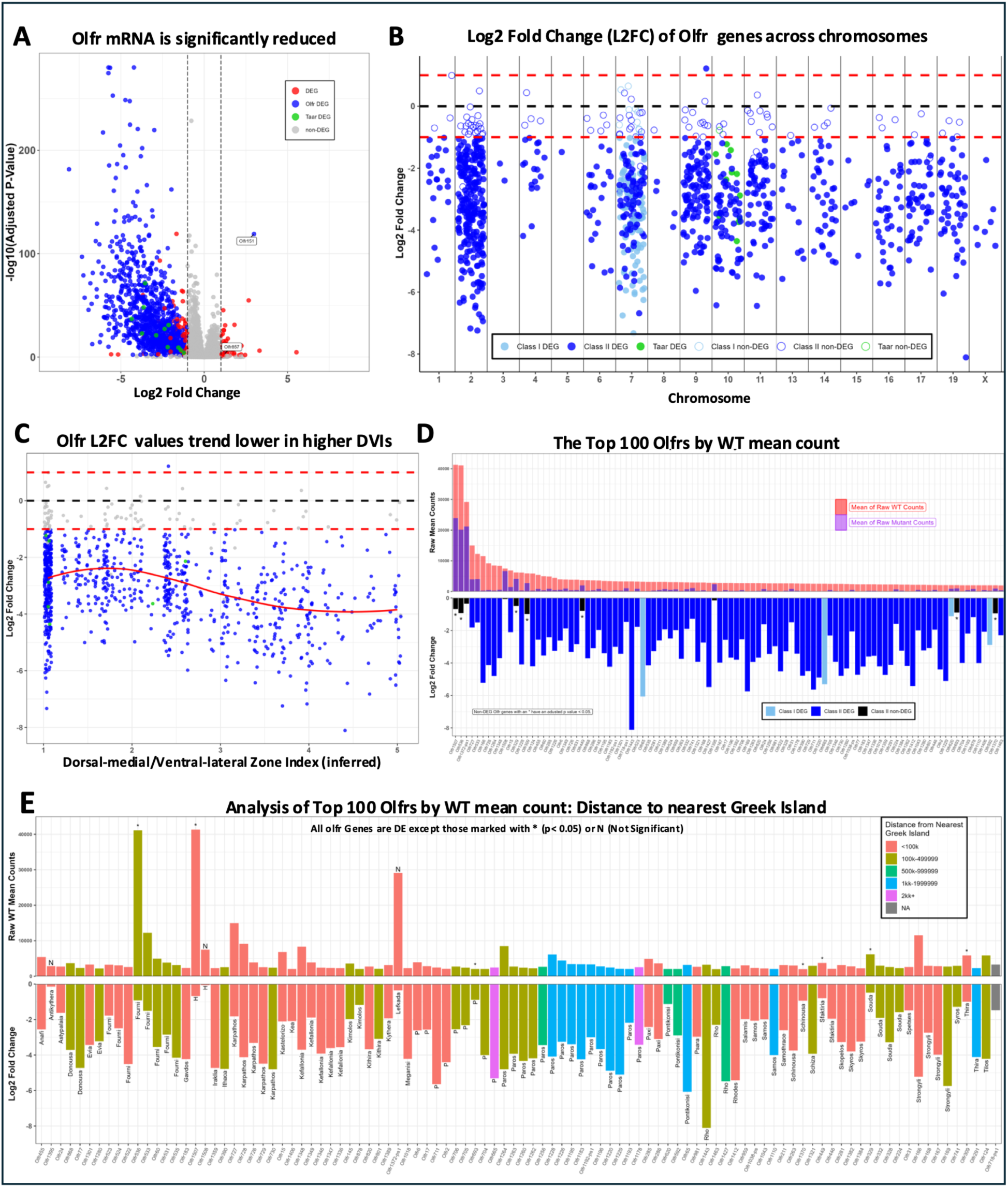
*OR10G4* expression is associated with significant loss of expression for nearly all OR (Olfr) genes across chromosomes, Dorsal-Ventral Index values, and Counts. (**A**). Volcano plot showing Log2 Fold Change for all genes categorized by DEG status (Differentially Expressed Gene). DEG set to a combination of <= −1 L2FC or >= 1 L2FC (black dashed lines) and p-value <= 0.05. Olfr151 shares sequences with the OR10G4 transgene, thus its elevated expression. (**B**). OR Log2 Fold Change split by chromosome and Class (I/II/TAAR) with the same DEG parameters as (**A**). (**C**). OR Log2 Fold Change separated by Dorsal Ventral Index (DVI) associated with each OR. Not all OR genes are represented due to “low expression” or “unusual” DVIs. DEG parameters consistent with (**A**). (**D**). Top 100 OR (Olfr) genes by mean of raw counts from WT samples. Top Panel, overlay of mean raw counts from WT and OR10G4 samples. Bottom Panel, Log2 Fold Change value for each OR, colored by Class and DEG status. *represents p <= 0.05, but not DEG as per (**A**). (**E**). The same top 100 OR (Olfr) genes by mean raw counts from WT samples, colored by proximity to the nearest Greek Island (GI) in *cis*. Top Panel. Mean raw count from WT. As in (D), * represents p <= 0.05, but not DEG. N is Not Significant. Bottom Panel. L2FC labeled with the name of the most proximal GI.

**Figure 3:**
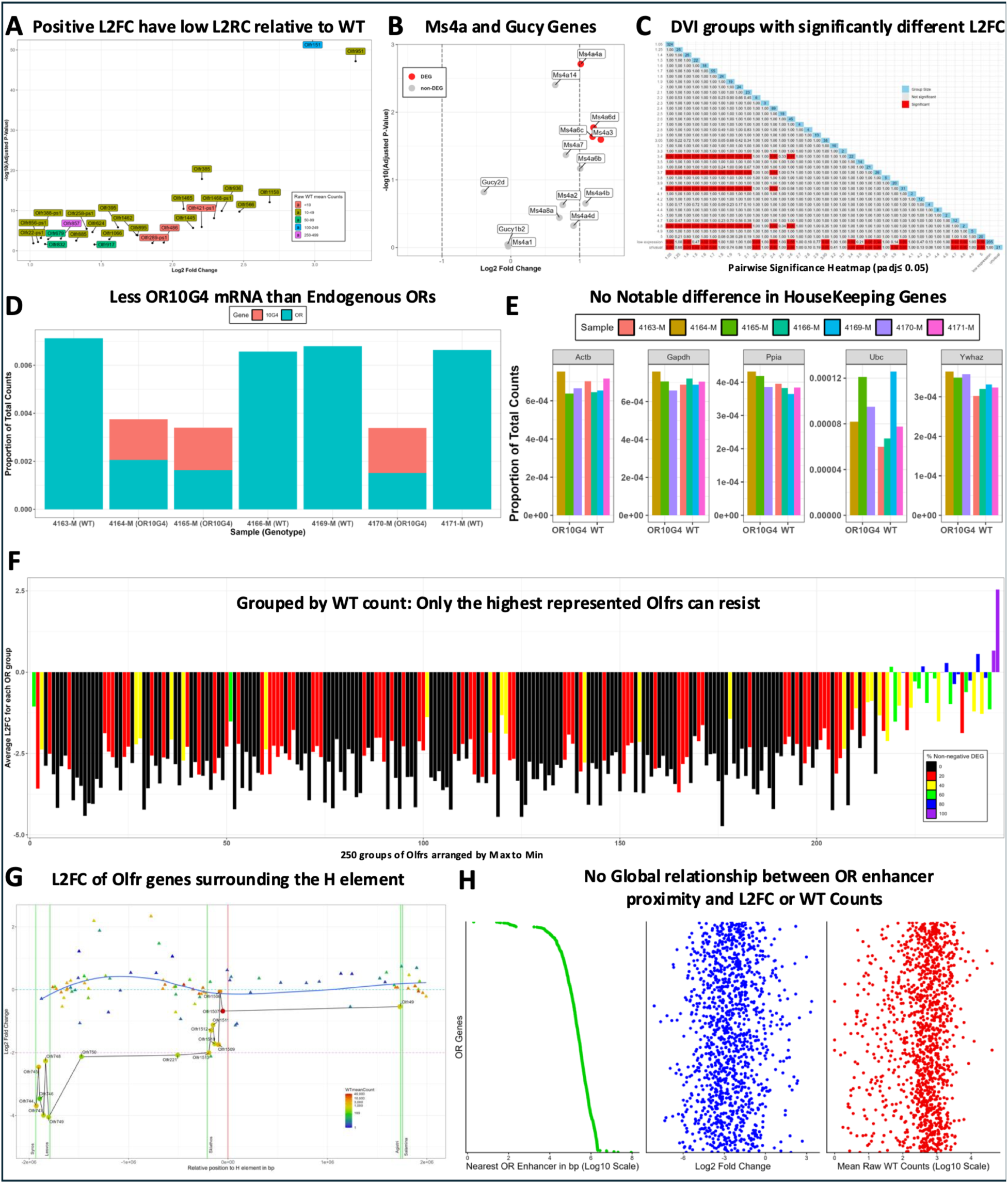
*OR10G4* suppresses nearly the entire OR repertoire and Greek Islands provide effectively no refuge for local OR genes. (**A**). Volcano plot showing Log2 Fold Change (L2FC) for non-negative DEG OR genes. DEG set to combination of <= −1.0 L2FC or >= 1.0 L2FC and p value <= 0.05. Colored by Mean raw count from WT samples. (**B**). Volcano plot showing L2FC for *Ms4a* and *Gucy* genes. DEG defined as in (A). (**C**). Pairwise plot highlighting DVI groups with significantly different L2FC values. Multiple comparisons P values adjusted with Holm correction; significance set to 0.05. (**D**). Comparing the normalized total endogenous OR and OR10G4 mRNA in each sample. Normalized to sample total RNA-seq counts. (**E**). Counts for each sample show similar values for presumptive housekeeping genes. Normalized to sample total RNA-seq counts. (**F**). OR genes were grouped together to evaluate mean L2FC and non-negative DEG across the spectrum of counts. OR genes with very low counts are least consistent, likely an artifact of sequencing rare transcripts from the entire repertoire. (**G**). Locus map for the H element, covering all OR and non-OR genes +/-2.25 million base pairs of H, with L2FC on the y axis and colors representing mean raw count from WT. H element centered and indicated with a red line. Other GI positions indicated with green lines. Circles are ORs and triangles are non-OR genes. Blue curve shows L2FC trend for non-OR genes. (**H**). An alignment of proximity to the nearest OR Enhancer (or Greek Island) for each OR (Left), the associated L2FC (Middle), and the mean raw count from WT (Right).

If *OR10G4* expression suppresses endogenous receptors via the native OR gene choice mechanism, one might expect loss of endogenous *OR* transcripts to be quantitatively offset by increased OR10G4 expression. However, this was not the case. OR10G4 transcript levels in OR10G4#7 neurons are relatively low per cell compared to OR10G4#10 neurons, as shown by RNAscope analyses (**Figure S1E**). After normalizing bulk RNA-seq counts to library size, we found that endogenous OR transcripts were reduced by approximately 75% in OR10G4#7 mice, whereas *OR10G4* expression restored total OR transcript abundance to only ∼50% of wild-type levels, leaving a significant overall reduction in steady-state OR mRNA abundance (p = 8.8e-0.6, **Figure 3D**). Single-cell RNA-seq analyses later confirmed this observation. Importantly, housekeeping gene expression remained stable between genotypes; for example, *Ywhaz* expression was barely elevated in OR10G4#7 samples (Holm-corrected p = 0.022). However, this difference could reflect sequencing depth or library quality, neither of which can explain the reduced steady-state OR transcript levels (**Figure 3E**).

OR transcript abundance reflects the number of OSNs expressing a given *OR* gene or the probability of accessing the singular choice machinery(6). As such, highly expressed ORs might be expected to resist suppression by *OR10G4* better. Indeed, among the 25 most highly expressed *OR* genes in wild-type animals, seven were resistant to transcript reduction, and this number increased to only ten among the top 100 *OR* genes (**Figure 2D**). However, when ORs were ranked by expression level and grouped into bins of five genes, suppressed receptors were distributed throughout the entire expression spectrum with no discernible trend (**Figure 3F**).

Although the three most abundant receptors (*Olfr1507*, *Olfr536*, and *Olfr1372-ps1*) were relatively resistant to suppression, this relationship did not extend to the broader *OR* repertoire. As expected, the few *OR*s exhibiting positive fold changes were among the lowest-abundance transcripts and likely reflect noise associated with low-count RNA-seq measurements (**Figure 3A**). One Greek Island enhancer, the H enhancer element from which the ×21 enhancer was derived, regulates distance-dependent expression of *Olfr1507*, *Olfr1508*, and *Olfr1509* (*24, 25*). Two of these genes are among the ten most highly expressed ORs and were largely resistant to suppression by *OR10G4* expression. While our data support the previously observed relationship between enhancer proximity and expression level, the expression changes observed for this H-dependent cluster were not readily explained by a simple competition model between the endogenous H enhancer and the *9×21-OR10G4* transgene (**Figure 3G**). Furthermore, while non-OR genes did not show a distance-dependent relationship with expression changes, genes near the H enhancer element tended to show increased expression, suggesting that Greek Islands can influence local non-OR transcriptional activity in the absence of their functioning as OR enhancers.

Finally, neither among the top 100 OR genes nor across the entire repertoire did we detect a meaningful relationship between proximity to any other Greek Island enhancers and either OR gene expression level or sensitivity to OR10G4-mediated suppression (**Figures 2E** and **3H**). Collectively, these observations argue that most Greek Islands do not function as classical strong enhancers controlling OR transcriptional output. Instead, their considerable distance from neighboring OR promoters and the widespread suppression caused by OR10G4#7 suggest that Greek Islands are unlikely to regulate steady-state OR expression levels or provide substantial competitive resistance against the 9×21 enhancer.

### OR10G4#7 scRNA-seq clustering, lineage analysis, and exploration of singularity

To determine how *OR10G4* expression affects singular OR gene expression, we performed single-cell RNA sequencing (scRNA-seq) of olfactory epithelium from wild-type and OR10G4 mice. Standard processing identified all expected olfactory epithelial cell populations (**Figure 4A, S3 and S4**). Although *Mbnl1* expression was broadly detected throughout the OR10G4 sample, *OR10G4* expression was restricted to *Omp*-positive cells of the olfactory sensory neuron (OSN) lineage, indicating that integration within the *Mbnl1* locus did not confer an *Mbnl1*-like expression pattern (**Figure S3F**).

**Figure 4.**
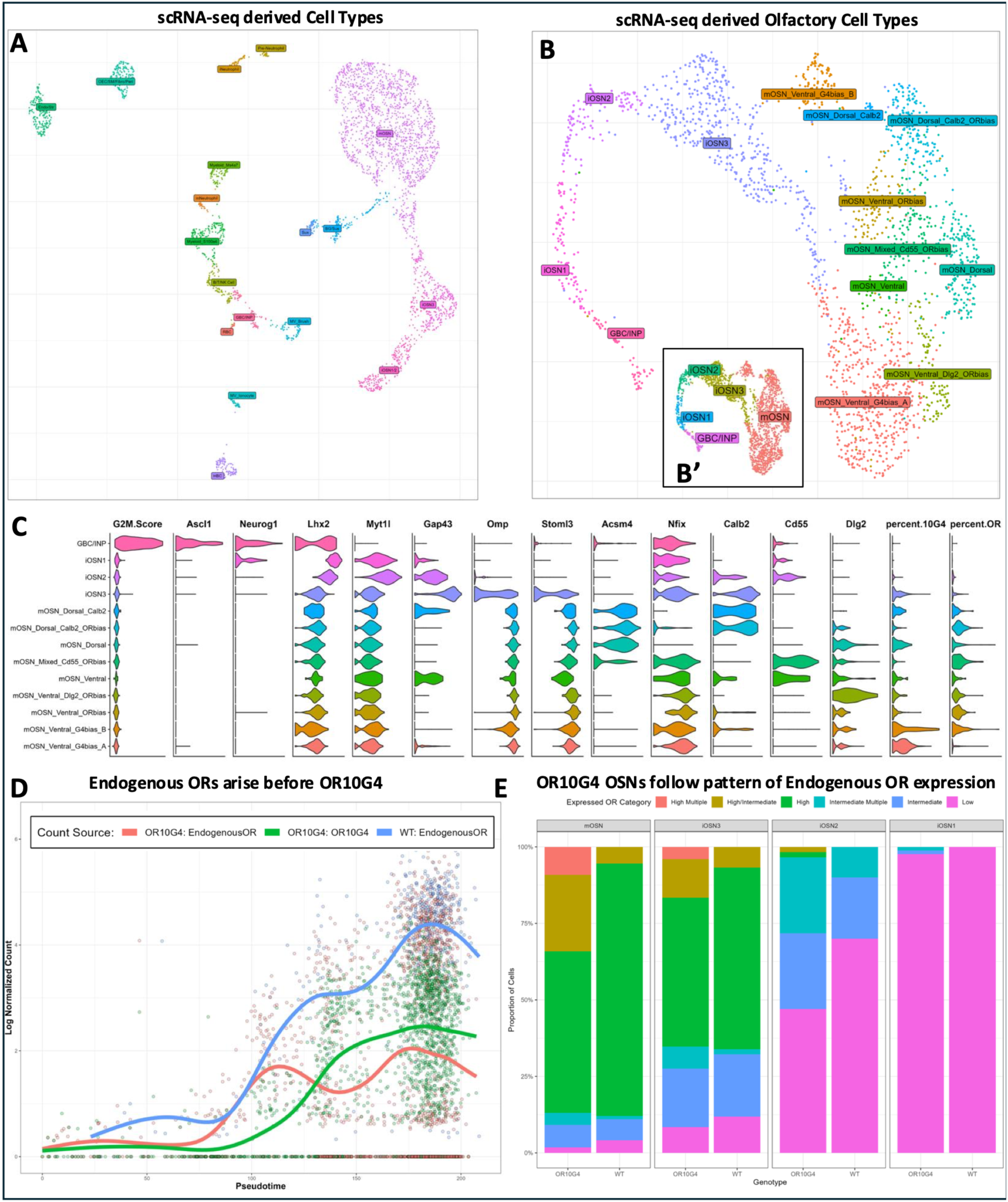
OR10G4 mice generate all expected cell types of the Nasal Epithelium but deviates OSN development away from Singular OR expression. (**A**). UMAP presentation of OR10G4 & WT cells distributed across known and expected Nasal cell types. Immature OSNs (iOSNs) and mature OSNs (mOSNs) are comprised of constituent clusters whose labels were aggregated for simplicity. (**B**). UMAP presentation of cell subset focusing on the GBC-to-mOSN lineage. OR genes removed prior to clustering. Selected cluster resolution produces several groups of mOSNs, further labeled based on dorsal/ventral markers, OR/OR10G4 expression, and/or other genes. (B’) Inset. Simplified cluster labels consolidate the mOSN clusters. (**C**). Gene expression of several OSN Lineage markers that define specific cell types in the lineage and explain labels in (B). (**D**). Pseudotime alignment with log-normalized counts for endogenous OR genes from WT and OR10G4 samples, compared to OR10G4 expression, to evaluate the relative timing of initiating gene expression. (**E**). Proportion barplot comparing OR expression state per OSN between Genotypes across four developmental stages, immature OSNs 1-3 (iOSN1-3) and mature OSNs (mOSN). “Low” labels are dismissed in the context of non-Low expression and only the top 2 OR expression categories are evaluated when applying labels to individual OSNs.

After identifying major cell classes, we reanalyzed only OSN-lineage cells and removed all *OR* genes, including *OR10G4*, from the clustering process to avoid receptor-driven clustering artifacts (**Figures S5A-5D**). OSN-lineage clusters were assigned using established marker genes and grouped into immature populations (GBC, INP, iOSN1-3) and mature OSNs (mOSNs) (**Figures 4B**, **4C, and S6**). While *Mbnl1* expression was relatively uniform across the lineage (see also mRNA and protein, **Figures S2A-2B**), *OR10G4* expression varied substantially among clusters, suggesting additional layers of regulation beyond transgene insertion site (**Figure S5E**).

Although *OR10G4* was detected in 94.9% of mOSNs, its expression levels were substantially lower than those of endogenous *OR* genes. Compared with endogenous receptors, *OR10G4* exhibited a 6.8-fold lower maximal normalized count and a 3.9-fold lower median expression level among OR-expressing mature OSNs (**Figure S7A**). Across all expressed genes, *OR10G4* ranked 473rd of 18,443 genes, and 410th of 583 ORs (**Figure S7B**), confirming that its dominance in the epithelium is not driven by unusually high transcript abundance.

Pseudotime analysis further revealed that endogenous *OR* expression commenced during late iOSN1 development and increased steadily into mature OSNs. In contrast, *OR10G4* expression was first detected during early iOSN2 and plateaued during the iOSN3-to-mOSN transition (**Figures 4D and S7C**). Endogenous OR expression in OR10G4 mice initially followed a pattern like wild type but diverged as *OR10G4* expression increased, suggesting that OR10G4 acts after endogenous OR choice has begun and exerts its effects during later stages of OSN maturation.

To assess singular OR gene expression, we categorized OR transcripts as High, Intermediate, or Low abundance. We excluded low-level transcripts from receptor-identity assignment. As expected, iOSN1 cells contained only low-level *OR* expression. In wild-type mice, cells progressively transitioned toward singular intermediate- or high-level *OR* expression as they matured. In contrast, OR10G4 mice maintained substantial *OR* co-expression through iOSN2, iOSN3, and mOSN stages (**Figure 4E).** This increase in co-expression was almost entirely attributable to OR10G4-positive neurons, as endogenous OR-only populations progressed similarly in the two genotypes (**Figure S7D**).

Using these criteria, 93.2% of wild-type mature OSNs were classified as effectively singular (**Table S2**). In contrast, *OR10G4* was detected in 94.9% of mOSNs, with 33.5% of cells co-expressing *OR10G4* and at least one endogenous *OR*, perhaps during Post-Selection Refinement (26) or as an example of mature OSNs failing to enforce singular *OR* gene expression. High-level *OR10G4*-expressing neurons were effectively singular in 77.8% of cases (WT High: 93.8%), whereas only 20.1% of intermediate-level *OR10G4* cells met the same criterion (WT Intermediate: 51.9%). Overall, *OR10G4* reproduced singular expression in 62.9% of mature OSNs while remaining co-expressed with endogenous *OR*s in a substantial fraction of the population, consistent with either incomplete post-selection refinement or reduced enforcement of singularity in mature neurons.

To test whether OR10G4 expression influences host-cell identity, we compared transcriptional states using UMAP embeddings generated without *OR* genes. Consistent with previous studies, mature OSNs expressing the same endogenous *OR* clustered together in transcriptional space (**Figure S8A**). *OR10G4*-expressing cells were distributed broadly across the UMAP, indicating that *OR10G4* can be expressed in multiple OSN subtypes. Nevertheless, co-expression of *OR10G4* frequently shifted endogenous OR-associated transcriptional profiles toward those of singular *OR10G4*-positive neurons (**Figures S8B-S8D**). Thus, *OR10G4* not only reflects existing OSN identities but can also influence the transcriptional state of its host neuron.

We further examined these relationships using consensus non-negative matrix factorization (cNMF). We identified 24 gene-expression programs (GEPs), 18 of which were deemed meaningful, including dorsal- and ventral-associated programs marked by *Acsm4* and *Nfix*, respectively (**Figure S9B**). After identifying candidate dorsal and ventral GEPs, we generated a dorsal-ventral (DV) score as previously described (**Table S4**, (27, 28)), allowing individual cells, clusters, and *OR*-specific mOSN populations to be positioned along the dorsal-ventral axis (**Figure S9C-S9E**). Singular OR10G4-expressing mOSNs exhibited a DV score within 0.125 of zero (**Figure S9E**), consistent with an intermediate dorsal-ventral identity or, alternatively, partial disruption of canonical dorsal-ventral transcriptional programs by *OR10G4* expression. To parallel the UMAP-distance analyses, we examined the 100 *OR* populations observed both in singular expression states and during co-expression with *OR10G4* for changes in their DV scores (**Figure S9F**).

*OR10G4*-expressing neurons occupied an intermediate dorsal-ventral position and, when co-expressed with endogenous *OR*s, shifted the associated neuronal populations toward this OR10G4-like state (**Figure S9F**). We next identified three GEPs that positively correlated with *OR10G4* expression but were only weakly correlated with one another. Combining these programs generated a composite score enriched in *OR10G4*-positive cells (**Figure S9G**). Because the constituent GEPs were associated with stress- and stress-adjacent transcriptional programs, we termed this metric the Tension Score.

The Tension Score was elevated in singular *OR10G4*-expressing mOSNs and increased in most endogenous *OR*-positive populations when *OR10G4* was co-expressed. The strongest effects were observed in *OR* populations exhibiting the lowest baseline Tension Scores (**Figures S9H-S9I**). Among the 100 *OR*-associated mOSN populations examined, 57 showed increases greater than 10%, whereas 61 showed some increase relative to their singular-expression baseline. Notably, 29 of the 33 *OR* populations with the lowest baseline Tension Scores showed elevated scores after *OR10G4* co-expression, suggesting that *OR10G4* exerts its greatest influence on transcriptionally less stressed neuronal populations. Together, these analyses show that *OR10G4* is expressed in most mature OSNs despite relatively modest transcript abundance, frequently coexists with endogenous *OR*s, and measurably reshapes the transcriptional state of the neurons in which it is expressed. *OR10G4* co-expression consistently shifts mOSNs toward an OR10G4-like molecular identity.

### Proteomic Analysis of Olfactory Cilia Reveals Dominance of OR10G4 Protein

Olfactory cilia constitute the primary interface between odorants and the olfactory system and contain a high concentration of odorant receptor proteins in OSNs(29). We performed quantitative mass spectrometry on cilia isolated from the olfactory epithelium of wild-type and OR10G4#7 mice to confirm OR10G4 protein presence and quantify remaining endogenous OR levels. Overall proteome quality was highly similar between genotypes. Samples displayed comparable peptide count distributions and total signal intensity, detected approximately 85%–90% of all unique peptide species, and shared ∼66% of identified peptides (**Figures S10A-S10C**). Consistent with our bulk RNA-seq results, all OR10G4 samples contained substantially less endogenous OR protein than wild-type controls (**Figure 5A**). Tryptic fragments and LC-MS of OR10G4 can only yield three non-hydrophobic small fragments (YLAISYPLR, YTSMMSGSR, MLMTLVSPSGR), all of which were detected and specific to OR10G4#7 cilia, confirming successful transcript translation. In contrast to the relatively low steady-state *OR10G4* mRNA levels observed by RNA-seq, OR10G4 protein expression nearly compensated for the reduction in endogenous OR proteins, effectively filling much of the receptor-protein deficit (**Figure 5B**).

**Figure 5:**
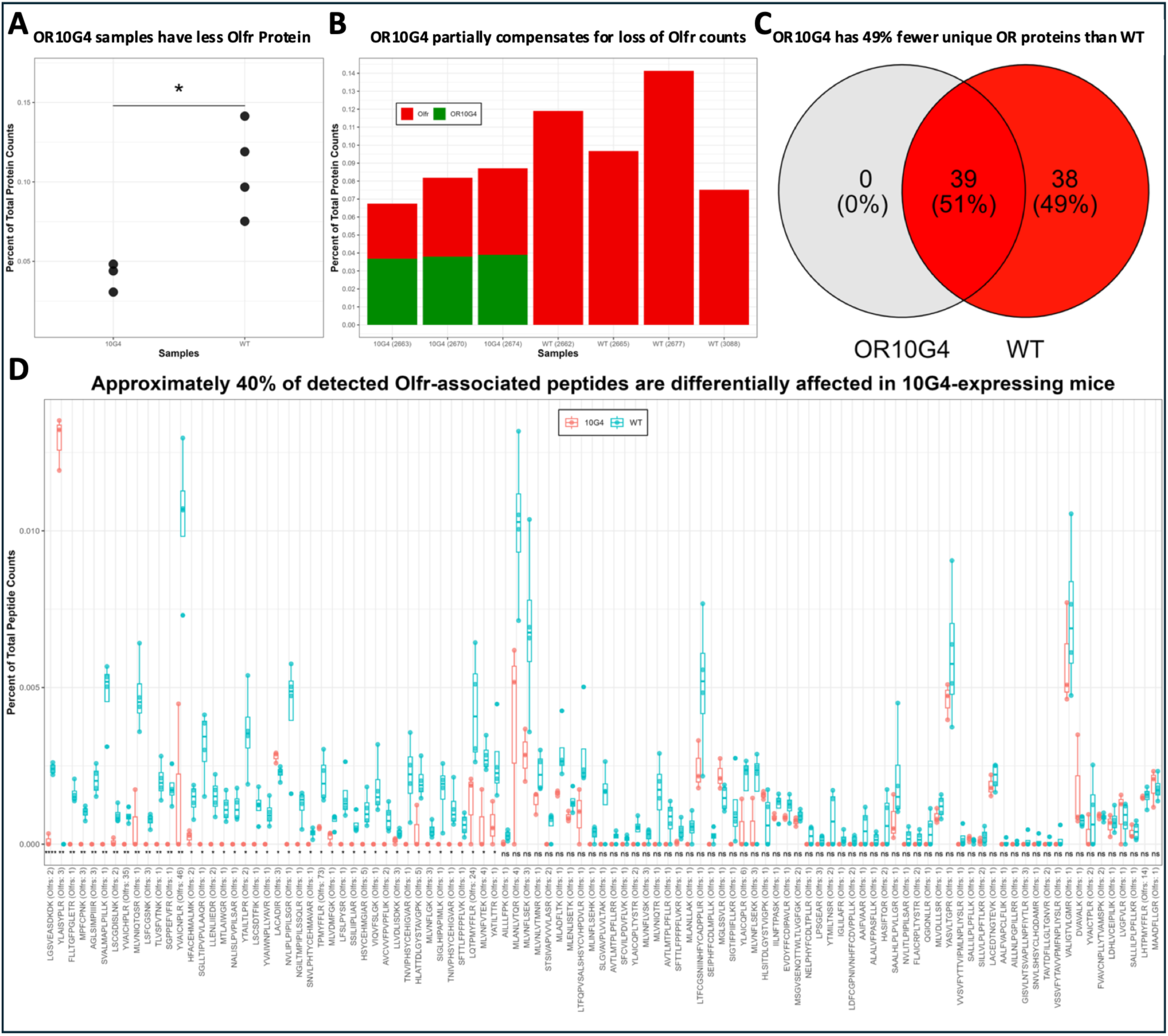
OR10G4 expression is associated with loss of Endogenous OR protein. (**A**). Comparison of OR (Olfr) protein counts for both genotypes, normalized by total protein counts per sample. * represents P value <= 0.05. (**B**). Sample-specific OR (Olfr) and/or OR10G4 protein counts, normalized by total protein counts per sample. (**C**). Venn Diagram showing the number of OR species detected in each genotype. Default peptide count aggregation method for identifying proteins selected for this comparison. 62 protein groups identified accounting for 77 OR species. (**D**). OR (Olfr) peptide-specific comparison between genotypes. Counts normalized to individual sample totals and presented as percent of total. Detected peptide sequences and the number of matching OR proteins on the X axis. No corrections for multiple comparisons applied to t-test values. Significance: *p <= 0.05, ** p <= 0.01, ***p <= 0.001, ****p <= 0.0001, ns = Not Significant.

Because our analyses were restricted to cilia isolated from the dorsal olfactory epithelium, we detected only a subset of the full OR repertoire. In this context, we also detected every OR protein identified in OR10G4 samples in wild-type samples. However, only 39 of the 77 OR proteins detected in wild-type animals were detected in OR10G4 mice (**Figure 5C**). To ensure that this result was not an artifact of peptide-to-protein aggregation, we repeated the analysis at the peptide level and observed a similar pattern (**Figure S10D**). Of the 103 OR-derived peptides detected across all samples, 58 (∼56%) were absent from OR10G4 mice. Because several peptides were inconsistently detected among individual wild-type replicates, only 42 of these peptides (∼41%) reached statistical significance (**Figure 5D**). Among the 44 peptides detected in both genotypes, approximately 30% were significantly altered, with several additional peptides showing similar trends (**Figure S10E**). The widespread absence of endogenous OR peptides in OR10G4 mice suggests that most detectable OR proteins in the dissected epithelial region were successfully sampled and that OR10G4 expression substantially reduces or eliminates a large fraction of the endogenous OR proteome. Thus, although *OR10G4* transcripts accumulate at lower steady-state levels than many endogenous *OR* mRNAs, OR10G4 protein abundance approaches endogenous receptor levels while endogenous OR proteins are markedly depleted. The presence of substantial levels of endogenous OR proteins suggests that the observed coexpression of *OR10G4* in scRNA-seq is also being observed in the proteomics data.

These findings have an important practical consequence: the OR10G4 transgenic line generates an olfactory cilia preparation with greatly reduced endogenous OR background and correspondingly enhanced OR10G4 signal. This improved signal-to-noise ratio enables accurate characterization of the OR10G4 ligand repertoire and provides a powerful platform for linking receptor activation to odor perception.

### Olfactory cilia bioassay with guaiacol derivatives correlated to human psychophysics data

Cilia-based Evaluation of Ligands and receptor InterActions (CELIA) isolated from our transgenic mice was used to characterize human OR1A1, OR5AN1, and OR10G4. In vitro expression of OR10G4 identified guaiacol, vanillin, and ethyl vanillin as agonists, raising the possibility that OR10G4 contributes to perception of the broadly defined “vanilla-like” odor quality(21). To determine whether OR10G4 is a primary receptor for vanilla-like odorants, we profiled a diverse panel of compounds that included (I) guaiacol-related molecules lacking a vanilla-like odor character, (II) chemically diverse odorants described as vanilla-like, and (III) a series of halogenated and alkylated guaiacol derivatives for which human perceptual-threshold data are available (**Tables S5**;(22, 23)).

Surprisingly, many odorants classified as vanilla-like failed to significantly activate OR10G4 above wild-type controls (**Figure 6A**). Similarly, most odorants lacking guaiacol-associated descriptors such as vanilla-like, smoky, or ham-like failed to activate OR10G4. The notable exception was 2-ethoxyphenol, a close structural analog of guaiacol that differs only by replacement of the methoxy substituent with an ethoxy group. Among the halogenated and alkylated guaiacol derivatives, approximately two-thirds were reported to possess vanilla-like qualities. However, 4-iodoguaiacol, despite its vanilla-like descriptor, failed to significantly activate OR10G4. In contrast, the strongest agonist identified in our screen was 5-bromoguaiacol, which is not typically described as vanilla-like. Collectively, these data argue against OR10G4 serving as a primary detector of vanilla odor quality. These data are consistent with OR10G3 having a 10-fold lower EC50 activation for ethyl vanillin than OR10G4(30).

**Figure 6:**
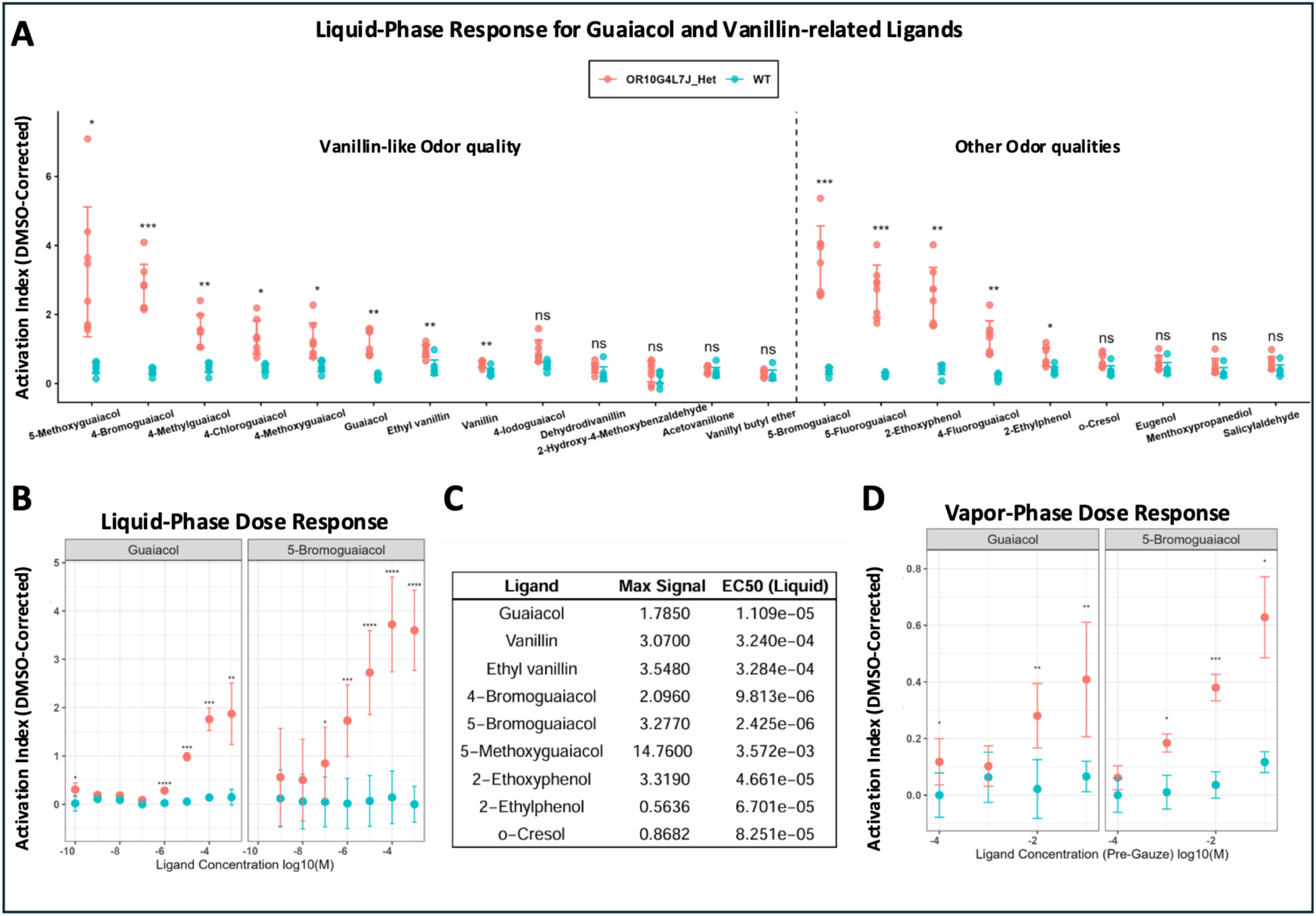
Activation of OR10G4 by Guaiacol derivates (**A**). WT and OR10G4-derived cilia tested with a panel of odors characterized with or without a “Vanilla-like” odor quality to identify if a preference exists. cAMP competition assay results were corrected using DMSO responses. Each ligand was tested 3+ times, and multiple-comparisons-corrected p-values were averaged on the log10 scale. Individual experimental values are plotted. (**B**). Ligand Dose Response shows that 5-Bromoguaiacol generates higher Activation Index values than Guaiacol at most significant dose values. Each ligand was tested 3 times in triplicate, and the average Activation Index was plotted. No p-value corrections were applied. (**C**). Summary table for Liquid Dose Response experiments. (**D**). Vapor Dose-Response results mimic Liquid Dose-Response results and validate *in vitro* measurements. Each ligand was tested once in triplicate at each dose. No p-value corrections were applied. Error bars: Activation Index mean +/-sd. Significance: * p <= 0.05, ** p <= 0.01, *** p <= 0.001, **** p <= 0.0001, ns = Not Significant.

To validate the superior activity of 5-bromoguaiacol, we performed dose-response analyses using selected odorants (**Figures 6B**, **6C, and S11**). Among all compounds tested, 5-bromoguaiacol exhibited the lowest EC50 and the greatest apparent potency. Although 5-methoxyguaiacol achieved a higher fitted maximal response, the large variance in its measurements produced an unreliable curve fit that extended beyond physiologically meaningful concentrations. Thus, 5-bromoguaiacol emerged as the most robust low EC50 OR10G4 ligand.

We next compared receptor activity with human olfactory thresholds. For compounds with published perceptual threshold values, vapor-phase detection thresholds correlated strongly with OR10G4 activation indices (**Figure 7A**), supporting the use of liquid-phase cAMP measurements to identify biologically relevant low EC50 ligands. To confirm these findings under more natural stimulus conditions, we developed a vapor-phase activation assay in which we exposed cilia preparations to odorants delivered through the headspace of sealed vials. Consistent with the liquid-phase experiments, 5-bromoguaiacol elicited stronger activation of OR10G4 than guaiacol across a range of vapor concentrations (**Figure 6D**).

**Figure 7:**
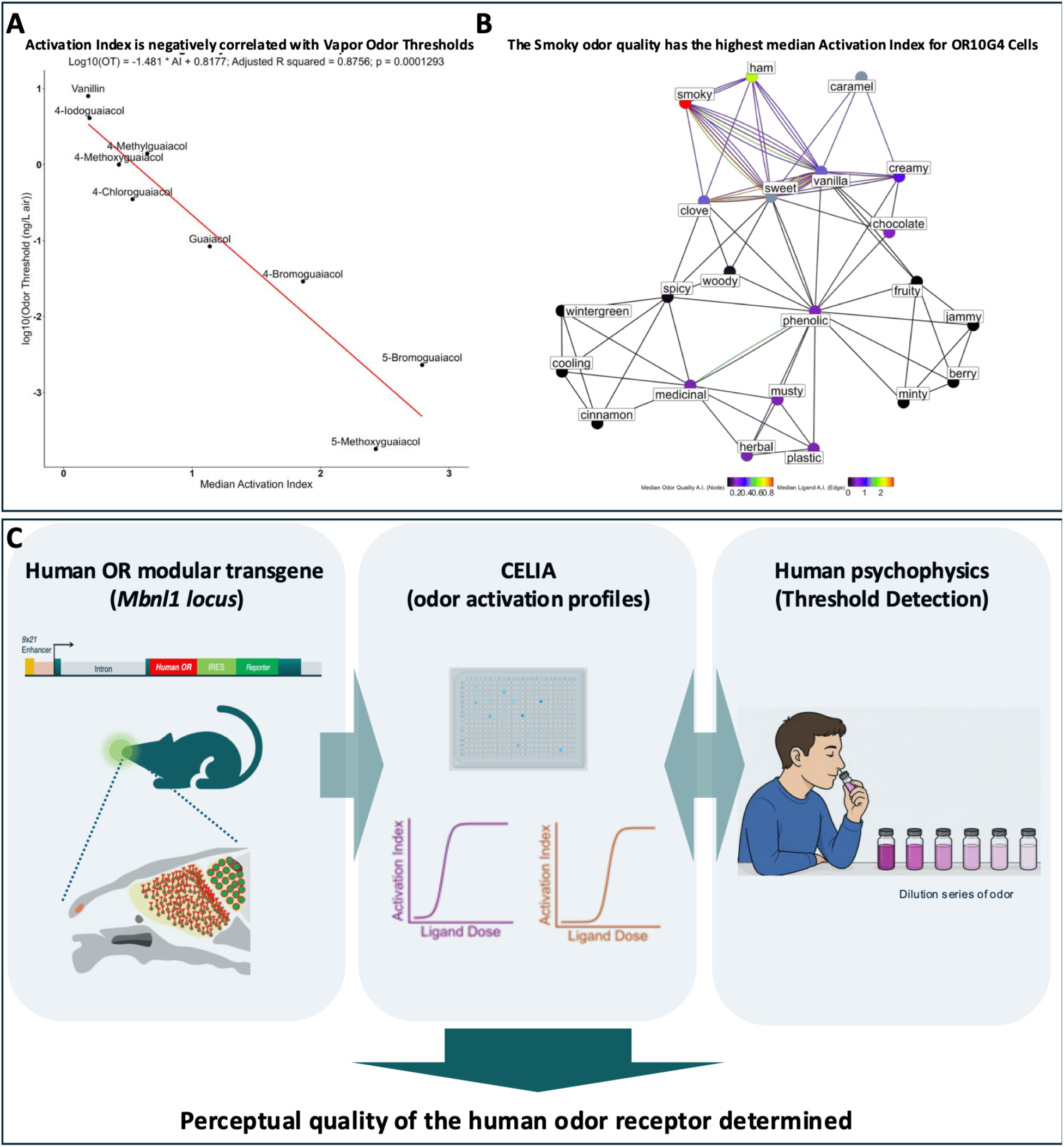
OR10G4-containing cilia are preferentially activated by odors defined by “smoky ham” qualities over the “vanilla” quality. (**A**). Human participant-based odor thresholds and experimentally derived Activation Index values for tested odors are strongly correlated, further validating our cAMP assay results. (**B**). Network analysis shows that the “Smoky Ham” nodes are associated with the highest activation index values. Individual lines represent odors that have odor qualities of the two connected nodes; a single odor can form many such lines depending on its collection of odor qualities. The color of a line represents the median A.I. value for that odor molecule. Nodes are colored based on the median of A.I. values associated with all lines that connect to that node. (**C**). Application of the OR10G4 transgenic platform for odor receptor profiling and psychophysical validation. (Left) Schematic of the human OR transgene cassette containing a human odorant receptor (hOR) coding sequence, an internal ribosome entry site (IRES), and a fluorescent reporter under the control of ×21-enhanced Olfr151 regulatory elements to drive expression in large populations of olfactory sensory neurons (OSNs). (Middle) CELIA (Cilia-based Evaluation of Ligands and receptor InterActions) assay of isolated olfactory cilia are screened against odorant dilution series using multiwell microtiter plates to quantify receptor-dependent signaling responses. (Right) A human psychophysical odor-threshold testing paradigm is used to determine responses across decreasing odor concentrations and compare them with receptor activation profiles measured in CELIA. This platform enables direct comparison of odorant receptor activation and human perceptual sensitivity, providing a framework for linking individual odorant receptors to specific odor-quality dimensions.

To determine whether OR10G4 is associated with a specific perceptual odor dimension, we constructed an odor-quality network linking tested odorants to their reported descriptors and colored network nodes according to median OR10G4 Activation Index values. This analysis revealed that the smoky odor category was associated with the strongest OR10G4 activation, followed closely by descriptors related to ham-like aromas (**Figure 7B**). In contrast, vanilla-like odorants displayed substantially weaker and less consistent activation patterns. Notably, the human detection threshold for 5-bromoguaiacol is lower than that for 4-bromoguaiacol, matching our activation data. Thus, 5-bromoguaiacol has the highest activation of any odor tested, and the lowest thresholds. More broadly, these findings show that combining receptor profiling with psychophysical data at odor-detection thresholds can directly link individual human odorant receptors to specific perceptual odor dimensions (**Figure 7C**).

## Discussion

### Why is 9×21 Expression at the Mbnl1 Locus Special?

We previously demonstrated that multimerization of the 21-bp homeodomain enhancer markedly increases the number of OSNs expressing OR transgenes, as observed in both the *5×21-OR1A1* and *5×21-OR5AN1* lines(10, 11). We subsequently mapped the *5×21-OR1A1* transgene to chromosome 1, far from any endogenous OR locus(4), indicating that elevated OR expression does not require integration within an OR cluster. Among the lines we have maintained, OR10G4#7 exhibits the most extensive OR gene expression identified to date. Importantly, however, widespread OR gene expression is not unique to this line. We also observed similarly broad expression in a *9×21-OR* founder, although we did not propagate that line. These observations suggest that the genomic environment capable of supporting high-frequency OR choice is not unique to the OR10G4#7 integration site and may occur at multiple locations throughout the genome.

One explanation is that highly active *n×21* transgenes preferentially achieve dominance when inserted into regions of chromatin that are already accessible in the olfactory sensory neuron lineage. Consistent with this possibility, the OR10G4#7 transgene is integrated within the *Mbnl1* locus, a region that displays strong ATAC-seq accessibility in OSNs. The combination of a powerful choice-promoting enhancer array (*9×21*) and a permissive chromatin environment may substantially increase the likelihood that the transgene engages the OR gene-choice machinery. In this model, the transgene is either more likely to recruit, or be recruited by, the molecular factors that govern singular OR gene selection. Taken together, these findings suggest that genomic position alone does not determine widespread OR choice.

### The Cis Regulation of OR Gene Choice

We identified a single-copy OR minigene (∼5 kb) consisting of a *9×21* enhancer, the *Olfr151* regulatory elements (promoter, first exon, and first intron), and a second exon containing the coding sequence of the human odorant receptor *OR10G4*. This transgene integrated randomly into the *Mbnl1* locus and acquired adjacent genomic sequences to serve as its 3′ untranslated region. Functional analyses indicate that the principal regulatory element governing expression of this minigene is the *9×21* enhancer(13). Remarkably, the OR10G4#7 insertion generates an almost monoclonal olfactory epithelium, with most olfactory sensory neurons selecting OR10G4.

This phenotype is fundamentally different from previously described artificial systems that drive widespread OR expression using OSN-specific *tTA* drivers (*Omp*, *Gng8*, or *CamKII*) paired with *TetO*-regulated OR transgenes(16, 17, 26, 31). In those systems, the tTA protein contains a potent viral transactivation domain that likely utilizes or bypasses endogenous mechanisms governing OR choice. By contrast, OR10G4#7 relies exclusively on native OR regulatory sequences and a cis-acting enhancer, providing a unique opportunity to study receptor selection within a more physiological framework.

The OR10G4#7 transgene is notable because it functions independently of known OR super-enhancers, termed Greek Islands (GIs), which are thought to play a central role in singular OR expression. Current models propose that approximately ∼180 Greek Islands in a diploid OSN aggregate into a small number of multi-enhancer assemblies, known as GI hubs, and that competition among thousands of cis enhancers within OR promoters for access to these hubs drives singular OR gene choice. Greek Islands are defined by overlapping LHX2 and Olf1/Ebf1 (O/E) ChIP-seq peaks within regions enriched for H3K9me3 chromatin and are present in nearly all mature OSNs. Interestingly, the *Olfr151* promoter itself contains a paired LHX2-O/E motif(13), an additional LHX2 site, and a separate O/E site, all within an H3K9me3-enriched region. However, because ChIP does not show uniform LHX2 and Olf1/Ebf1 occupancy across all OSNs, we do not classify the Olfr151 promoter as a Greek Island. The nearest bona fide Greek Island, **Kimolos,** is located approximately 23 kb away. Importantly, the Olfr151 minigene functions efficiently as an independent transgene(7, 8, 13), suggesting that proximity to **Kimolos** is not required for receptor activation.

Approximately 90 Greek Islands putative enhancers have been identified in the haploid OR subgenome, corresponding to roughly 180 sites in a diploid OSN(19, 32). ATAC-seq profiling of the OSN lineage shows these regions are constitutively accessible. Yet the 189-bp *9×21* enhancer promotes expression of its linked OR in most OSNs while effectively outcompeting the endogenous OR repertoire. This observation echoes earlier studies showing that placement of the H enhancer adjacent to an *Olfr1507* minigene resulted in expression throughout approximately 80% of OSNs(33). Although both produce strikingly similar phenotypes, the 9×21 sequence lacks paired LHX2-O/E sites and would not be classified as a Greek Island enhancer.

Notably, both transgenes outperform not only endogenous receptors linked to the same enhancers but also the entire complement of endogenous Greek Islands, including both endogenous H elements. In each case, the enhancer is positioned close to the OR promoter: approximately 5 kb for *H-Olfr1507* and ∼500 bp for *9×21-OR10G4*. By contrast, experimental multimerization of six Greek Islands (**Lipsi**, **Sfaktiria**, **Crete**, H, and two **Rhodes** elements) to preassemble a synthetic GI hub resulted in only a modest, approximately three-fold increase in expression of the neighboring receptors *Olfr1413* and *Olfr1414*, which lie ∼27 kb away on either side of the engineered locus(19).

It is difficult to reconcile the relatively modest activity of this six-GI synthetic hub with the dramatic expression achieved by endogenous H-linked ORs, the *H-Olfr1507* transgene, or the *9×21-OR10G4* transgene. These observations suggest that enhancer proximity and intrinsic enhancer strength may be more important determinants of OR gene choice than assembling large multi-enhancer hubs. The preferential expression of *H-Olfr1507* and *9×21-OR10G4* relative to a synthetic six-GI hub argues that local cis-regulatory interactions possess greater functional potency than GI hubs alone. Rather than serving primarily as transcriptional amplifiers, Greek Islands may instead increase the probability that an OR locus engages the gene-choice machinery. In contrast, strong local enhancers such as H element and *9×21* appear capable of dominating this process, allowing their linked ORs to achieve expression frequencies far exceeding those of endogenous receptors. These findings support a model in which cis-regulatory strength, enhancer proximity, and chromatin accessibility are the principal determinants of OR choice probability.

### Competition Between 9×21-OR10G4 and Greek Islands for Singular OR Gene Choice

Deletion of the H enhancer results in loss of expression of three of the eight linked OR genes: *Olfr1507* (∼52 kb away), *Olfr1508* (∼76 kb away), and *Olfr1509* (∼88 kb away). This result was explained by H having both cis and trans activating properties. In contrast, simultaneous deletion of three Greek Islands (**Lipsi**, **Sfaktiria**, and H) primarily affected genes neighboring **Lipsi** and **Sfaktiria** without completely abolishing their expression. These data were interpreted as evidence that the remaining ∼184 Greek Islands could still assemble functional GI hubs capable of rescuing OR gene expression. The implications are that endogenous *OR* expression reflects a probability distribution, with each *OR* having a distinct likelihood of being activated by the available pool of GI hubs, except in the case of the H enhancer.

Bulk RNA-seq provides a useful measure of these probabilities because OR transcript abundance correlates closely with the number of OSNs expressing a given receptor. By this metric, *Olfr1507* is the most highly expressed OR gene in the genome and is associated with the largest OSN population. *Olfr1508* ranks approximately tenth among all ORs, whereas the more distal *Olfr1509* ranks near the top 100. In the OR10G4#7 background, however, these genes respond very differently. *Olfr1507* remains the most highly expressed OR despite a 42% reduction in transcript abundance, *Olfr1508* decreases by only 10% and remains among the highest-expressed receptors, whereas *Olfr1509* declines by approximately 75%, like more than half of the endogenous OR repertoire. One possible explanation for the apparent resilience of *Olfr1508* is that it is frequently coexpressed with another OR in the OR10G4#7 background. Indeed, scRNA-seq identified four *Olfr1508*-positive neurons, all of which coexpressed a second OR gene. Nevertheless, the highly non-linear effects observed across the H-dependent cluster—42% reduction for *Olfr1507*, 10% for Olfr1508, and 75% for *Olfr1509*—argue against a simple model in which the *9×21-OR10G4* transgene passively interferes with a fixed pool of 5–6 GI hubs assembled from ∼190 endogenous Greek Islands.

More generally, little evidence suggests that the OR10G4 transgene acts by sequestering LHX2 or GI-associated factors. Previous studies identified more than 16,000 LHX2-bound sites throughout the genome, making it unlikely that adding only nine LHX2-binding sites within the *9×21* enhancer substantially alters global LHX2 occupancy. Furthermore, if Greek Islands were the primary determinants of OR gene expression, OR genes located near strong or abundant GI elements would be predicted to show increased resistance to suppression by *OR10G4*. We observed no such relationship. Across nearly 1,000 OR genes, changes in expression were not correlated with local Greek Island proximity, nor did enhancer density predict resistance to suppression by expression of *OR10G4*. The lack of a relationship between OR expression changes and Greek Island proximity suggests that “OR10G4 effects” are not mediated through disruption of GI hubs. Instead, the data are more consistent with altered competition among local enhancer-promoter units for access to the OR choice machinery. In this model, OR expression is determined by the relative probability that a given enhancer-promoter pair successfully engages the mechanisms governing singular receptor selection.

We therefore propose that endogenous Greek Islands and GI hubs function as relatively weak regulatory elements that promote stochastic receptor choice rather than strong transcriptional activation. Such weak enhancers would be advantageous because they permit a broad OR repertoire to be represented across the olfactory epithelium. By contrast, the emergence of a particularly strong enhancer would bias OR gene choice toward a single OR and diminish receptor diversity. Such dominant regulatory elements would likely be selected against during evolution or constrained by genomic architecture. The H enhancer may represent an example of this principle, residing more than 50 kb from *Olfr1507*, a distance that could limit its effective strength despite its demonstrated importance for receptor expression. Our observations support a model in which singular OR gene choice is governed primarily by competition among enhancer-promoter units of differing strengths, whereas Greek Islands act as relatively weak regulatory elements that preserve the stochastic and diverse nature of endogenous receptor selection.

### Singular OR Gene Choice: First Come, First Served or Winners and Losers?

In the prevailing GI Hub model of singular OR gene expression, early competition among multiple OR loci produce transient co-expression states from which a single “winning” receptor ultimately emerges. Under this framework, any OR that gains an early transcriptional advantage should be favored for selection. However, our previous experiments showed that driving *Olfr151* expression from the early O/E2 promoter at the iOSN1 stage does not alter either the singular or stochastic nature of OR choice, suggesting that early transcription alone is insufficient to determine receptor selection(34).

Likewise, if exogeneous *OR10G4* is simply competing with endogenous ORs for access to GI hubs, then we might expect more transcription per cell. We previously showed that the increased number of *Olfr151*-expressing neurons in *4×21-Olfr151* mice does not correlate with increased OR mRNA abundance per neuron, nor does the expanded expressing population alter Olfr151 axonal convergence within its native cell type (4, 10). Consistent with these observations, OR10G4#7-expressing neurons exhibit lower steady-state transcript levels than OR10G4#10 neurons by RNAscope analysis (**Figure S1E**), and single-cell RNA-seq reveals that *OR10G4* transcript abundance in OR10G4#7 is, on average, two- to four-fold lower than that of endogenous OR genes (**Figure S7A**). This reduction likely reflects transgene truncation and use of a de novo 3′UTR acquired at the integration site.

These findings further decouple OR gene choice from OR transcript abundance. If receptor selection were determined primarily by mRNA levels, *OR10G4* in OR10G4#7 would not be expected to dominate the olfactory epithelium despite producing fewer transcripts per neuron than many endogenous ORs. Instead, our data suggest that OR choice and OR transcription are distinct processes. The *9×21* transgene appears to use the endogenous OR gene choice machinery while accessing it at substantially higher frequencies than endogenous OR loci.

### Reconciling the Mechanism of OR Gene Choice

We propose a model in which LHX2 binds accessible target sites throughout the genome and nucleates an LDB1-mediated aggregation process whereby proximally or distally located dimers of LHX2 are bound by dimers of LDB1. Greek Islands are particularly high-probability LHX2 binding sites because LHX2 occupies them in nearly all OSNs. This privileged status may derive from the close juxtaposition of LHX2 and Olf1/Ebf1 (O/E) motifs together with the presence of multiple LHX2 binding sites within individual enhancers. For example, the H enhancer contains four potential LHX2 binding sites, one positioned adjacent to an O/E motif. This framework accommodates many observations interpreted through the GI Hub model while introducing a critical distinction: neither LHX2 binding sites nor GI hubs themselves establish transcriptional singularity. Instead, we propose that they facilitate recruitment to a separate singular nuclear body where OR transcription occurs. In this model, our 189-bp *9×21* enhancer, despite lacking O/E sites, remains highly accessible to LHX2 and is particularly effective at finding this transcriptional compartment. Once associated with the nuclear body, OR transcription is stabilized and singular expression emerges.

Under this view, neither the 21-bp homeodomain element nor endogenous Greek Islands function as rheostats that regulate OR transcription magnitude. Rather, they act as targeting elements that determine the probability that an OR locus gains access to the singular transcriptional compartment. Our single-cell sequencing data support this interpretation, as most OSNs express one dominant receptor throughout OR gene choice. Occasional low-level expression of secondary ORs likely reflects transient or unstable interactions with the nuclear body rather than true violations of singular expression. Such low-level transcription would be unlikely to influence axon guidance or odor coding. In this sense, OR gene choice is best described as pseudo-singular: one receptor overwhelmingly dominates transcriptional output, while rare secondary transcriptional events persist at biologically insignificant levels.

A key prediction of this model is that a highly efficient enhancer should dramatically suppress endogenous receptor expression by preferentially recruiting its linked OR locus to the singular nuclear body. The OR10G4#7 line supports this prediction. Single-cell RNA sequencing revealed an approximately 75% reduction in expression across *Class I OR*s, *Class II OR*s, and *TAAR* genes. We and others previously proposed that all three receptor classes utilize a common LHX2-dependent gene-choice mechanism. In the OR10G4#7 background, nearly all endogenous receptor genes are substantially downregulated, including many of the most highly expressed ORs. These observations support a unified model of receptor selection and indicate that OR gene choice operates through fundamentally similar mechanisms across *Class I OR*, *Class II OR*, and *TAAR*-expressing neurons(4). Our findings support a model in which singular OR expression arises through competition among receptor loci for access to a solitary transcriptional compartment.

### Odor Responses Measured from Isolated Cilia

OR10G4 was previously identified as a receptor for guaiacol (2-methoxyphenol), an aromatic compound containing adjacent hydroxyl and methoxy substituents on a benzene ring. Adding an aldehyde group at the 5-position yields vanillin, while replacing the methoxy group with an ethoxy group yields ethyl vanillin. Although OR10G4 responds to these compounds, aldehyde substitution substantially increases receptor EC50 relative to guaiacol, suggesting that OR10G4 is not optimized for detecting vanilla odorants. However, it remained formally possible that OR10G4 represented the principal, or even sole, receptor responsible for vanilla perception in humans.

To evaluate this possibility, we tested OR10G4 against a comprehensive panel of odorants commonly described as possessing a vanilla-like quality. Several canonical vanilla odorants failed to activate OR10G4, indicating that activation of this receptor is neither necessary nor sufficient for perception of vanilla odor. Although odor transport, metabolism, or partitioning in the nasal mucus may alter receptor access to some ligands, our data suggest that OR10G4 is unlikely to be the primary receptor mediating vanilla perception. We also observed no fundamental differences between vapor-phase and liquid-phase odor delivery in our receptor activation assays(11).

Having largely excluded a central role for OR10G4 in encoding vanilla odor quality, we next asked whether OR10G4 instead defines a perceptual space associated with guaiacol derivatives. Two independent psychophysical studies demonstrated that substitutions at the 5-position of guaiacol substantially reduce human detection thresholds relative to corresponding substitutions at the 4-position. Halogen substitutions (bromo-, chloro-, and iodo-) and alkyl substitutions such as methoxy produced dramatic increases in odor potency. Participants most often described these compounds as smoky, sweet, or related odor qualities.

Strikingly, our receptor activation assays mirrored the psychophysical observations. Across multiple substitution classes, 5-position modifications consistently generated lower EC50 OR10G4 ligands than corresponding 4-position substitutions, and several of the strongest agonists identified in our study contained 5-position substitutions. This concordance between receptor activation and human detection thresholds suggests that OR10G4 contributes directly to the perceptual sensitivity of this odorant class.

Network analysis linking odorants to their associated perceptual descriptors further supported this interpretation. When odor descriptors were weighted by median OR10G4 activation, the strongest associations were observed for the smoky and, to a lesser extent, ham-like odor categories. These findings suggest that OR10G4 contributes to the receptor code underlying smoky odor perception in humans and may also help represent cured-meat or ham-like aromas. If individuals lose the ability to activate OR10G4, they can still smell 5-bromoguaiacol, likely through OR10G3, OR10G7, OR2J2, or OR7A17(30), but may lose the ability to distinguish between smoky odors and those OR-associated odor qualities (defined by EC50 and perceptual thresholds). In this regard, odor detection does not guarantee unambiguous perception.

More broadly, the OR10G4 study illustrates a general strategy for assigning odor qualities to individual human odorant receptors. By identifying receptor agonists across a range of EC50s and combining these data with psychophysical threshold measurements, it should be possible to map perceptual odor dimensions to specific human ORs. Our data are in line with the idea that each OR carries primacy for odor responses in an olfactory map with sparse activation (35, 36) as well as an associated perceptual quality(37), where ORs set the threshold of detection for a given odor (36, 38).

We are currently applying the *×21*-expression system to candidate receptors responsive to sebum-derived odorants associated with Parkinson’s disease (PD)(39). The OR10G4#7 locus provides a scalable platform for identifying receptors involved in PD-associated odor signatures and, more generally, for linking receptor activation profiles to human odor perception. Ultimately, this approach may enable receptor-level decoding of the human olfactory code.

## Conclusion

We propose that the OR10G4#7 mouse reveals a new framework for understanding singular OR gene choice, consistent with nearly two decades of observations showing that Greek Island (GI) regions of the olfactory subgenome aggregate near the expressed OR allele. Our model is rooted in our earliest observations that LHX2 binding sites are key determinants of aggregation into a solitary nuclear compartment where OR singularity is established. In contrast to the prevailing view that Greek Island hubs directly drive singular OR transcription, we suggest that a biochemically simpler nuclear body governs singular expression and serves as the site of OR activation. In this framework, cis-acting LHX2-bound sequences, including Greek Islands, promote recruitment of OR loci to this compartment, where singular transcription is established and maintained.

This model accommodates the well-documented aggregation of Greek Islands near active OR loci while assigning these elements a supporting rather than causative role in OR singularity. More broadly, such a mechanism may extend beyond the olfactory system. Monoallelic gene families and even X-chromosome inactivation may rely on analogous nuclear compartments that selectively activate one genomic locus while excluding others. In this view, X-chromosome inactivation could be reframed as preferential activation of a single X chromosome through association with a specialized nuclear body, whereas inactive X chromosomes fail to access that compartment.

At the circuit level, thousands of distinct ORs and naturally occurring OR variants direct axons to receptor-specific glomeruli. The transcriptional diversity associated with these receptors has made it difficult to disentangle the molecular mechanisms underlying singular gene choice from those governing OR-dependent axonal coalescence. The OR10G4#7 platform overcomes this limitation by combining a single-copy transgene insertion with a highly efficient choice-promoting enhancer, producing large populations of neurons expressing a common receptor. This system enables experimental approaches that move beyond transcriptional profiling and toward direct identification of the proteins and molecular interactions that mediate OR gene choice, homotypic axonal sorting, and glomerular formation.

At the same time, the near-monoclonal expression achieved in OR10G4#7 provides unprecedented access to receptor-specific cilia and neuronal populations, allowing detailed characterization of OR receptive ranges with greatly improved signal-to-noise ratios. By coupling these analyses with psychophysical data, we were able to associate OR10G4 with a specific perceptual odor-quality dimension. More generally, targeted replacement of the OR10G4 coding sequence with other human ORs will enable systematic mapping of receptor activation profiles to perceptual odor qualities. We therefore envision the OR10G4#7 locus as a scalable platform for linking OR gene choice, neuronal connectivity, receptor pharmacology, and odor perception, ultimately providing a framework for decoding the molecular logic of the human olfactory system (**Figure 7C**).

## Limitations of the Study

Although the data strongly suggest that the *9×21* enhancer increases the probability of OR choice, we cannot directly distinguish whether this occurs through increased recruitment to the OR gene choice machinery, increased retention within that machinery, and/or resistance to silencing. Similarly, while our findings are inconsistent with a simple GI-Hub competition model, they do not directly visualize *9×21* enhancer interactions with or without GI-Hub enhancers and the proposed nuclear body. Currently, we have no protein markers for the proposed nuclear body other than the transcription unit itself. Direct visualization of chromosomal interactions and nuclear architecture in the OR10G4#7 integrated locus will be required to distinguish between competing mechanistic models.

Our single-cell analyses also reveal extensive *OR10G4* co-expression with endogenous ORs. While these observations support a model of incomplete post-selection refinement or reduced enforcement of singularity, scRNA-seq captures only a snapshot of transcriptional state and cannot determine whether co-expression is stable or transient. RNA fluorescent *in situ* hybridization on OSNs will ultimately resolve whether transcription is singular or if cells maintain multi-OR states(40).

The proteomic analyses are restricted to cilia isolated from the dorsal olfactory epithelium and therefore sample only a subset of the endogenous OR repertoire. As a result, the OR protein depletion measured here likely underestimates changes across the entire olfactory epithelium. More comprehensive proteomic analyses spanning both dorsal and ventral regions will be needed to fully quantify the effects of OR10G4 on the endogenous receptor landscape.

Finally, our assignment of a smoky odor-quality dimension to OR10G4 is based on correlations among receptor activation, odor detection thresholds, and published psychophysical descriptors. In vitro analysis of the entire OR repertoire has led to the idea that only a small number of human ORs (OR10G3, OR10G4, OR10G7, OR2J2, and OR7A17) are involved in detection of guaiacol derivatives; however, these activations may correlate with in vitro expression level rather than true ability to activate the OR. In this context, our assay provides a reproducible mechanism to express any OR. Future studies combining receptor odor profiles from the entire human OR repertoire with human psychophysics will be necessary to define the extent to which individual ORs dictate specific perceptual odor qualities.

## Acknowledgments

We thank the Hunter College Animal Facility Manager Barbara Wolin and Veterinarian Patricia Glennon for help in maintaining the transgenic colony, and Rada Norinsky at the Transgenic Core Facility at The Rockefeller University for generating transgenic founders. We thank the WCMC Genomic Core Facility for sequencing. Many thanks to Charlotte D’Hulst and Yukie Takabake for initial work on OR10G4 odor profiling. Many Thanks to Mary Slavinsky for animal colony maintenance, to Martina Pyrski for sharing RNAscope protocol for fixed tissues. We thank Takasago for providing the following Vanillin-like odors: Vanillyl butyl ether and Menthoxypropanediol.

## Author Contributions

P.F. and M.O. conceived the project. P.F., E.L., M.O., R.M. designed the experiments. P.F., E.L., M.O, and R.M. performed all experiments and analyzed the data. P.F., E.L., and M.O. wrote the manuscript with input from R.M.

## Methods

**See Star Methods**

### Mice

Mice used in this study were bred and maintained in the Laboratory Animal Facility of Hunter College, CUNY. The Hunter College IACUC approved all procedures. Animal care and procedures were in accordance with the Guide for the Care and Use of Laboratory Animals (NHHS Publication No. [NIH] 85-23). The Hunter College IACUC approved all mouse experimental protocols.

## References

1. Feinstein P, Bozza T, Rodriguez I, Vassalli A, Mombaerts P. Axon guidance of mouse olfactory sensory neurons by odorant receptors and the beta2 adrenergic receptor. Cell. 2004;117(6):833–46. Epub 2004/06/10. doi: 10.1016/j.cell.2004.05.013. PubMed PMID: 15186782.

2. Feinstein P, Mombaerts P. A contextual model for axonal sorting into glomeruli in the mouse olfactory system. Cell. 2004;117(6):817–31. Epub 2004/06/10. doi: 10.1016/j.cell.2004.05.011. PubMed PMID: 15186781.

3. Bozza T, Feinstein P, Zheng C, Mombaerts P. Odorant receptor expression defines functional units in the mouse olfactory system. J Neurosci. 2002;22(8):3033–43. Epub 2002/04/12. doi: 20026321. PubMed PMID: 11943806; PMCID: PMC6757547.

4. Makhlouf M, D’Hulst C, Omura M, Rosa A, Mina R, Bernal-Garcia S, Lempert E, Saraiva LR, Feinstein P. A common mechanism of singular gene choice is revealed by broadly expressed odorant receptor transgenes within the olfactory epithelium. Cell Rep. 2025;44(7):115955. Epub 2025/07/15. doi: 10.1016/j.celrep.2025.115955. PubMed PMID: 40663455.

5. Bressel OC, Khan M, Mombaerts P. Linear correlation between the number of olfactory sensory neurons expressing a given mouse odorant receptor gene and the total volume of the corresponding glomeruli in the olfactory bulb. J Comp Neurol. 2016;524(1):199–209. Epub 2015/06/24. doi: 10.1002/cne.23835. PubMed PMID: 26100963; PMCID: PMC4758392.

6. Ibarra-Soria X, Levitin MO, Saraiva LR, Logan DW. The olfactory transcriptomes of mice. PLoS Genet. 2014;10(9):e1004593. Epub 2014/09/05. doi: 10.1371/journal.pgen.1004593. PubMed PMID: 25187969; PMCID: PMC4154679.

7. Rothman A, Feinstein P, Hirota J, Mombaerts P. The promoter of the mouse odorant receptor gene M71. Mol Cell Neurosci. 2005;28(3):535–46. Epub 2005/03/02. doi: 10.1016/j.mcn.2004.11.006. PubMed PMID: 15737743.

8. Vassalli A, Rothman A, Feinstein P, Zapotocky M, Mombaerts P. Minigenes impart odorant receptor-specific axon guidance in the olfactory bulb. Neuron. 2002;35(4):681–96. Epub 2002/08/27. doi: 10.1016/s0896-6273(02)00793-6. PubMed PMID: 12194868.

9. Vassalli A, Feinstein P, Mombaerts P. Homeodomain binding motifs modulate the probability of odorant receptor gene choice in transgenic mice. Mol Cell Neurosci. 2011;46(2):381–96. Epub 2010/11/30. doi: 10.1016/j.mcn.2010.11.001. PubMed PMID: 21111823; PMCID: PMC3746036.

10. D’Hulst C, Mina RB, Gershon Z, Jamet S, Cerullo A, Tomoiaga D, Bai L, Belluscio L, Rogers ME, Sirotin Y, Feinstein P. MouSensor: A Versatile Genetic Platform to Create Super Sniffer Mice for Studying Human Odor Coding. Cell Rep. 2016;16(4):1115–25. Epub 2016/07/12. doi: 10.1016/j.celrep.2016.06.047. PubMed PMID: 27396335.

11. Omura M, Takabatake Y, Lempert E, Benjamin-Hong S, D’Hulst C, Feinstein P. A genetic platform for functionally profiling odorant receptors in olfactory cilia ex vivo. Sci Signal. 2022;15(746):eabm6112. Epub 2022/08/10. doi: 10.1126/scisignal.abm6112. PubMed PMID: 35944068.

12. Shah A, Ratkowski M, Rosa A, Feinstein P, Bozza T. Olfactory expression of trace amine-associated receptors requires cooperative cis-acting enhancers. Nat Commun. 2021;12(1):3797. Epub 2021/06/20. doi: 10.1038/s41467-021-23824-3. PubMed PMID: 34145232; PMCID: PMC8213819.

13. Rosa A. The Seat of Singular Gene Choice: Dissecting the Role of Odorant Receptor Enhancer Elements That Lead to Mammalian Olfactory System Expression. CUNY Academic Works. 2024(Ph.D. Dissertation).

14. Cichy A, Dewan A, He Z, Fitzgerald C, Ratkowski M, Krasewicz J, Ozarkar V, Kaye S, Teng T, Zhang J, Feinstein P, Bozza T. A microbiome-derived olfactory signal regulates inter-male aggression and social dominance in mice. Curr Biol. 2026;36(8):1946–58 e4. Epub 2026/04/01. doi: 10.1016/j.cub.2026.03.009. PubMed PMID: 41916309; PMCID: PMC13094369.

15. Nguyen MQ, Zhou Z, Marks CA, Ryba NJ, Belluscio L. Prominent roles for odorant receptor coding sequences in allelic exclusion. Cell. 2007;131(5):1009–17. Epub 2007/11/30. doi: 10.1016/j.cell.2007.10.050. PubMed PMID: 18045541; PMCID: PMC2195930.

16. Nguyen MQ, Marks CA, Belluscio L, Ryba NJ. Early expression of odorant receptors distorts the olfactory circuitry. J Neurosci. 2010;30(27):9271–9. Epub 2010/07/09. doi: 10.1523/JNEUROSCI.1502-10.2010. PubMed PMID: 20610762; PMCID: PMC2906254.

17. Fleischmann A, Shykind BM, Sosulski DL, Franks KM, Glinka ME, Mei DF, Sun Y, Kirkland J, Mendelsohn M, Albers MW, Axel R. Mice with a “monoclonal nose”: perturbations in an olfactory map impair odor discrimination. Neuron. 2008;60(6):1068–81. Epub 2008/12/27. doi: 10.1016/j.neuron.2008.10.046. PubMed PMID: 19109912; PMCID: PMC2732586.

18. Monahan K, Horta A, Lomvardas S. LHX2- and LDB1-mediated trans interactions regulate olfactory receptor choice. Nature. 2019;565(7740):448–53. Epub 2019/01/11. doi: 10.1038/s41586-018-0845-0. PubMed PMID: 30626972; PMCID: PMC6436840.

19. Monahan K, Schieren I, Cheung J, Mumbey-Wafula A, Monuki ES, Lomvardas S. Cooperative interactions enable singular olfactory receptor expression in mouse olfactory neurons. Elife. 2017;6. Epub 2017/09/22. doi: 10.7554/eLife.28620. PubMed PMID: 28933695; PMCID: PMC5608512.

20. Markenscoff-Papadimitriou E, Allen WE, Colquitt BM, Goh T, Murphy KK, Monahan K, Mosley CP, Ahituv N, Lomvardas S. Enhancer interaction networks as a means for singular olfactory receptor expression. Cell. 2014;159(3):543–57. Epub 2014/11/25. doi: 10.1016/j.cell.2014.09.033. PubMed PMID: 25417106; PMCID: PMC4243057.

21. Mainland JD, Keller A, Li YR, Zhou T, Trimmer C, Snyder LL, Moberly AH, Adipietro KA, Liu WLL, Zhuang H, Zhan S, Lee SS, Lin A, Matsunami H. The missense of smell: functional variability in the human odorant receptor repertoire. Nature Neuroscience. 2014;17(1):114–20. doi: 10.1038/nn.3598 PMID - 24316890.

22. Juhlke F, Lorber K, Wagenstaller M, Buettner A. Influence of the Chemical Structure on Odor Qualities and Odor Thresholds of Halogenated Guaiacol-Derived Odorants. Front Chem. 2017;5:120. Epub 2018/01/13. doi: 10.3389/fchem.2017.00120. PubMed PMID: 29326924; PMCID: PMC5741668.

23. Schranz M, Lorber K, Klos K, Kerschbaumer J, Buettner A. Influence of the chemical structure on the odor qualities and odor thresholds of guaiacol-derived odorants, Part 1: Alkylated, alkenylated and methoxylated derivatives. Food Chem. 2017;232:808–19. Epub 2017/05/12. doi: 10.1016/j.foodchem.2017.04.070. PubMed PMID: 28490144.

24. Fuss SH, Omura M, Mombaerts P. Local and cis effects of the H element on expression of odorant receptor genes in mouse. Cell. 2007;130(2):373–84. Epub 2007/07/31. doi: 10.1016/j.cell.2007.06.023. PubMed PMID: 17662950.

25. Nishizumi H, Kumasaka K, Inoue N, Nakashima A, Sakano H. Deletion of the core-H region in mice abolishes the expression of three proximal odorant receptor genes in cis. Proc Natl Acad Sci U S A. 2007;104(50):20067–72. Epub 2007/12/14. doi: 10.1073/pnas.0706544105. PubMed PMID: 18077433; PMCID: PMC2148423.

26. Abdus-Saboor I, Al Nufal MJ, Agha MV, Ruinart de Brimont M, Fleischmann A, Shykind BM. An Expression Refinement Process Ensures Singular Odorant Receptor Gene Choice. Curr Biol. 2016;26(8):1083–90. Epub 2016/04/05. doi: 10.1016/j.cub.2016.02.039. PubMed PMID: 27040780.

27. Brann DH, Tsukahara T, Tau C, Kalloor D, Lubash R, Kannan LT, Klimpert N, Kollo M, Escamilla-Del-Arenal M, Bintu B, Schaefer A, Fleischmann A, Bozza T, Datta SR. A spatial code governs olfactory receptor choice and aligns sensory maps in the nose and brain. Cell. 2026;189(11):3358–79 e30. Epub 2026/04/30. doi: 10.1016/j.cell.2026.03.051. PubMed PMID: 42054991; PMCID: PMC13134484.

28. Tsukahara T, Brann DH, Pashkovski SL, Guitchounts G, Bozza T, Datta SR. A transcriptional rheostat couples past activity to future sensory responses. Cell. 2021;184(26):6326–43 e32. Epub 2021/12/09. doi: 10.1016/j.cell.2021.11.022. PubMed PMID: 34879231; PMCID: PMC8758202.

29. Li RC, Molday LL, Lin CC, Ren X, Fleischmann A, Molday RS, Yau KW. Low signaling efficiency from receptor to effector in olfactory transduction: A quantified ligand-triggered GPCR pathway. Proc Natl Acad Sci U S A. 2022;119(32):e2121225119. Epub 2022/08/02. doi: 10.1073/pnas.2121225119. PubMed PMID: 35914143; PMCID: PMC9371729.

30. Emter R, Merillat C, Buchli F, Flachsmann F, Natsch A. Decoding human olfaction by high heterologous expression of odorant receptors detecting signature odorants. Curr Biol. 2025;35(21):5252–63 e4. Epub 2025/10/12. doi: 10.1016/j.cub.2025.09.041. PubMed PMID: 41075782.

31. Fleischmann A, Abdus-Saboor I, Sayed A, Shykind B. Functional interrogation of an odorant receptor locus reveals multiple axes of transcriptional regulation. PLoS Biol. 2013;11(5):e1001568. Epub 2013/05/24. doi: 10.1371/journal.pbio.1001568. PubMed PMID: 23700388; PMCID: PMC3660300.

32. Wu H, Zhang J, Jian F, Chen JP, Zheng Y, Tan L, Sunney Xie X. Simultaneous single-cell three-dimensional genome and gene expression profiling uncovers dynamic enhancer connectivity underlying olfactory receptor choice. Nat Methods. 2024;21(6):974–82. Epub 2024/04/16. doi: 10.1038/s41592-024-02239-0. PubMed PMID: 38622459; PMCID: PMC11166570.

33. Serizawa S, Miyamichi K, Nakatani H, Suzuki M, Saito M, Yoshihara Y, Sakano H. Negative feedback regulation ensures the one receptor-one olfactory neuron rule in mouse. Science. 2003;302(5653):2088–94. Epub 2003/11/01. doi: 10.1126/science.1089122. PubMed PMID: 14593185.

34. Movahedi K, Grosmaitre X, Feinstein P. Odorant receptors can mediate axonal identity and gene choice via cAMP-independent mechanisms. Open Biol. 2016;6(7). Epub 2016/07/29. doi: 10.1098/rsob.160018. PubMed PMID: 27466441; PMCID: PMC4967819.

35. Giaffar H, Shuvaev S, Rinberg D, Koulakov AA. The primacy model and the structure of olfactory space. PLoS Comput Biol. 2024;20(9):e1012379. Epub 2024/09/10. doi: 10.1371/journal.pcbi.1012379. PubMed PMID: 39255274; PMCID: PMC11423968.

36. Burton SD, Brown A, Eiting TP, Youngstrom IA, Rust TC, Schmuker M, Wachowiak M. Mapping odorant sensitivities reveals a sparse but structured representation of olfactory chemical space by sensory input to the mouse olfactory bulb. Elife. 2022;11. Epub 2022/07/22. doi: 10.7554/eLife.80470. PubMed PMID: 35861321; PMCID: PMC9352350.

37. Raps DA, Pierce, G.M., Papiani, G., Wu, L, Arroyave, R, Brann, J.H., and Pfister, P. Evidence of elemental encoding at the olfactory periphery. bioRxiv. 2025;10.1101/2025.04.21.649882.

38. Dewan A, Cichy A, Zhang J, Miguel K, Feinstein P, Rinberg D, Bozza T. Single olfactory receptors set odor detection thresholds. Nat Commun. 2018;9(1):2887. Epub 2018/07/25. doi: 10.1038/s41467-018-05129-0. PubMed PMID: 30038239; PMCID: PMC6056506.

39. Trivedi DK, Sinclair E, Xu Y, Sarkar D, Walton-Doyle C, Liscio C, Banks P, Milne J, Silverdale M, Kunath T, Goodacre R, Barran P. Discovery of Volatile Biomarkers of Parkinson’s Disease from Sebum. Acs Central Sci. 2019;5(4):599–606. doi: 10.1021/acscentsci.8b00879.

40. Mina Raena B. Employing High Probability Gene Choice Elements to Understand Singular Odorant Receptor Expression. CUNY Academic Works https://academicworkscunyedu/gc_etds/4071. 2020.

