## Supplemental Figures for "Functional Profiling of a Human Odorant Receptor Using Olfactory Cilia Links its Activation to Odor Quality"

#### Figure S1

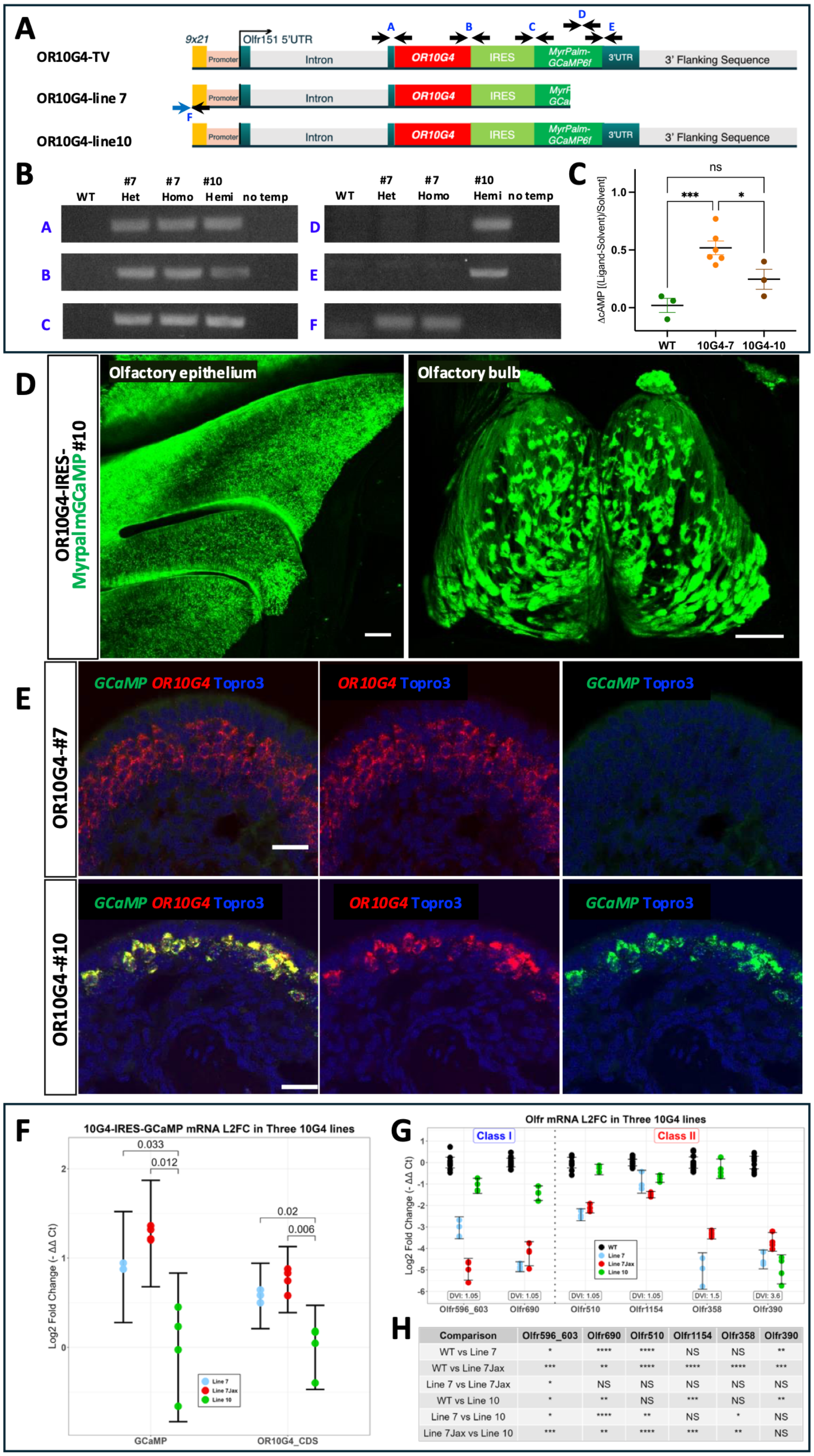


| Comparison | Olfr596_603 | Olfr690 | Olfr510 | Olfr1154 | Olfr358 | Olfr390 |
|---|---|---|---|---|---|---|
| WT vs Line 7 | * | **** | **** | NS | NS | ** |
| WT vs Line 7Jax | *** | ** | **** | **** | **** | *** |
| Line 7 vs Line 7Jax | * | NS | NS | NS | NS | NS |
| WT vs Line 10 | * | ** | NS | *** | NS | ** |
| Line 7 vs Line 10 | * | **** | ** | NS | * | NS |
| Line 7Jax vs Line 10 | *** | ** | **** | *** | ** | NS |

**Figure S1**, related to Figure 1: OR10G4 Line# 7 and line #10

(**A**). Schematic diagrams of the OR10G4 targeting vector and two OR10G4-expressing lines, line #7 and line #10. We analyzed the integrated transgene structures of two OR10G4-positive lines by genomic DNA PCR using 6 primer sets (**A-F** in blue). Primers shown with black arrows (11 primers) were in the targeting vector, and the forward primer of F (shown with a blue arrow) was in the *Mbnl1* locus, where the integration site of the truncated targeting vector in line #7 is located. (**B**). The results of genomic DNA PCR analysis of the structure of the integrated OR10G4 transgene in line #7 and line #10, with a wild-type (WT) control. Line #10 carries the whole targeting vector (positive for A-E), while line #7 carries OR10G4 but not the full-length GCaMP, and the amplicons in F show integration in the Mbnl1 locus. (**C**). Activation of extracted cilia samples from WT, line #7 (10G4-7), and line #10 (10G4-10) with the OR10G4 ligand, 50 μM guaiacol. Activation was measured by cAMP concentration after incubation with guaiacol or solvent (DMSO). Data are from three for WT and 10G4-10, and 6 for 10G4-7 biological replicates; error bars show the SEM. Data were analyzed by ordinary one -way ANOVA with Fisher's LSD test for each pair compared. ns $p=0.0768$, * $p< 0.05$ and ***$p<0.001$, (**D**). MyrPalmGCaMP6 expression in the MOE and olfactory bulbs of OR10G4 transgenic line #10. The left panel shows a medial whole-mount view of the main olfactory epithelium and the turbinate; scale bar 200 μm. The right panel shows a dorsal view of the olfactory bulbs of the line #10 mouse; scale bar: 500 μm. (**E**). RNAscope of coronal sections of the MOE from line #7 and line #10 with OR10G4 and GCaMP probes, counterstained with TOPRO3; scale bar 20 μm. (**F**). RT-qPCR targeting the marker and OR coding regions of *OR10G4* mRNA from three different OR10G4 lines. (**G**). RT-qPCR targeting OR coding regions of endogenous OR mRNA from three different OR10G4 lines. (**H**). Summary table for (G). For (F) & (G), 3-4 animals were tested per line, L2FC scaled relative to mean Line 10 or WT values, error bars are mean +/- 95% confidence interval, and p values were not corrected for multiple comparisons.

### Figure S2

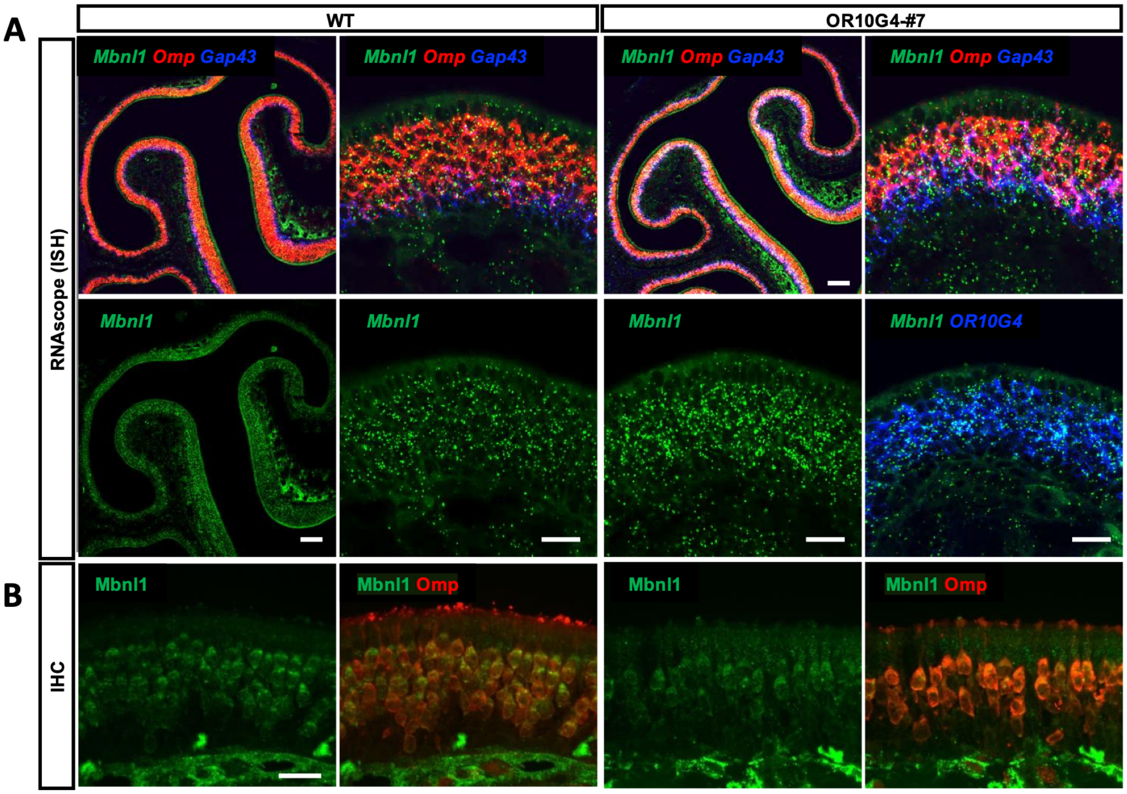
A
B
WT
OR10G4-#7
RNAscope (ISH)
IHC
Mbnl1 Omp Gap43
Mbnl1 Omp Gap43
Mbnl1 Omp Gap43
Mbnl1 Omp Gap43
Mbnl1
Mbnl1
Mbnl1
Mbnl1 OR10G4
Mbnl1
Mbnl1 Omp
Mbnl1
Mbnl1 Omp

**Figure S2**, related to Figure 1: *Mbnl1* expression in the olfactory epithelium

(**A**). RNAscope in the coronal section of the olfactory epithelium of wild-type (WT) and OR10G4-line#7 with *Mbnl1*, *Omp*, *Gap43* and *OR10G4* probes. *Mbnl1 is* expressed broadly in the olfactory epithelium in *Omp*- and *Gap43*-positive cells. Scale bar: low-magnification images 100 µm and high-magnification images 20 µm. (**B**). Immunohistochemistry on a coronal section of WT and OR10G4-line#7 with anti-MBNL1 antibody and anti-OMP antibody, scale bar 20 µm.

### Figure S3

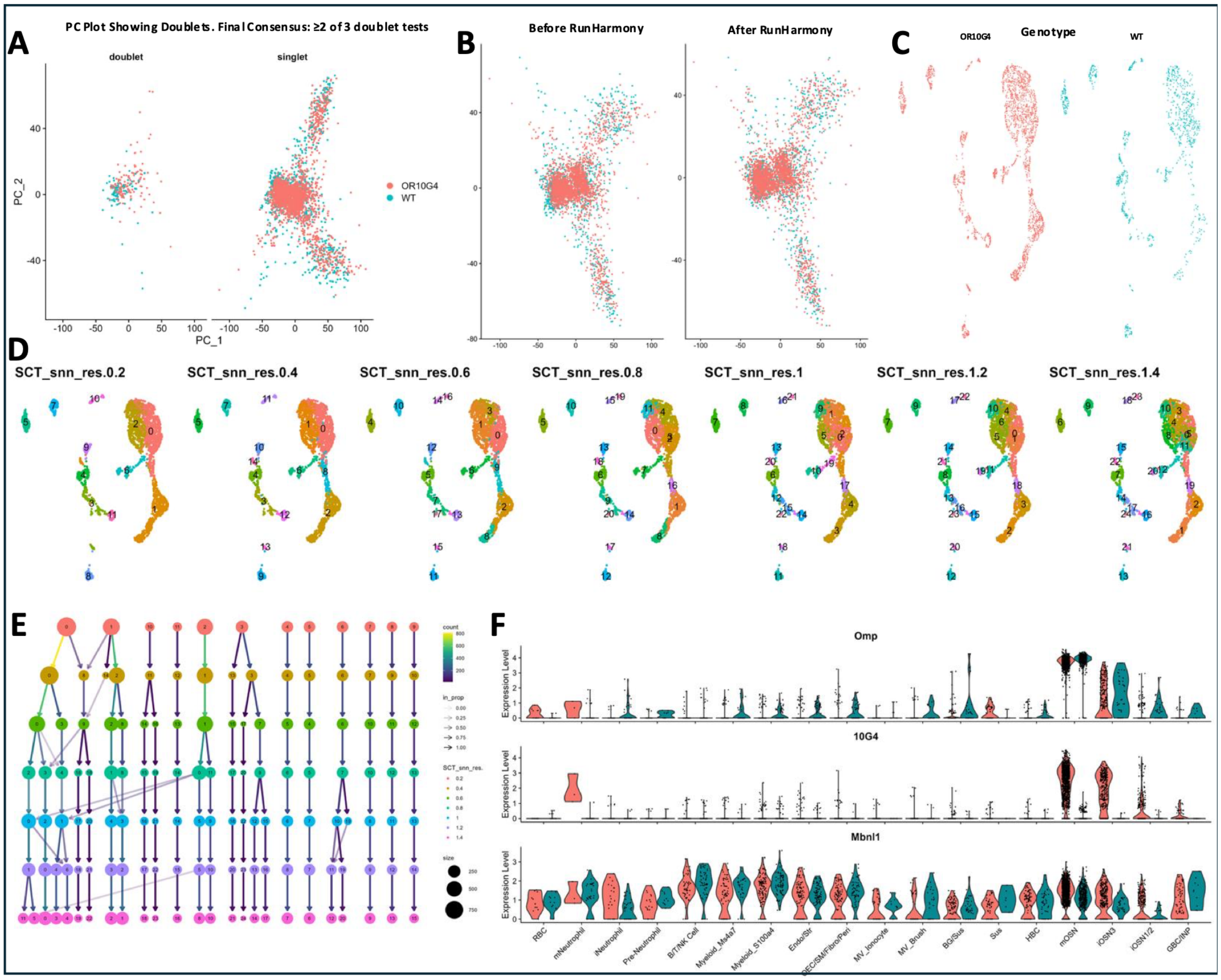
A
PC Plot Showing Doublets. Final Consensus: ≥2 of 3 doublet tests
doublet
singlet
PC_2
PC_1
OR10G4
WT
B
Before RunHarmony
After RunHarmony
C
Genotype
OR10G4
WT
D
SCT_snn_res.0.2
SCT_snn_res.0.4
SCT_snn_res.0.6
SCT_snn_res.0.8
SCT_snn_res.1
SCT_snn_res.1.2
SCT_snn_res.1.4
E
count
in_prop
SCT_snn_res.
size
F
Omp
10G4
Mbnl1
Expression Level
RBC
mNeutrophil
iNeutrophil
Pre-Neutrophil
B/T/NK Cell
Myeloid_Ms4a7
Myeloid_S100a4
Endo/Str
OEC/SM/Fibro/Peri
MV_Ionocyte
MV_Brush
BG/Sus
Sus
HBC
mOSN
iOSN3
iOSN1/2
GBC/INP

**Figure S3**, related to Figure 4: Combining and Clustering the OR10G4 and WT scRNA-seq datasets.

(**A**). Doublet analysis was used to identify likely doublets from the sequencing process, visualized across PCA space. Removed prior to additional clustering steps. (**B**). The Harmony package was used to better combine the two datasets, showing a subtle merging of genotype-specific groups across PCA space. (**C**). UMAP shows the contribution of OR10G4 and WT samples across the various cell clusters. Both genotypes contribute to all groups and thus all identified cell type categories. (**D**). Evaluating the impact of Resolution on total clusters via UMAP visualization. Lower resolution might group dissimilar cells, but greater resolution can create unnecessary splits. (**E**). The cluster tree helps select a reasonable resolution value. Simple cluster splits are easy to evaluate, but multi-cluster reshuffling suggests grouping instability. Highest resolution tested, 1.4, deemed adequate to account for all expected cell types (based on previous evaluation attempts). (**F**). Violin Plots with labeled Clusters looking at Omp expression as a measure of doublet presence and OR10G4 and Mbnl1 expression to look for blatant cross-influence on the expression profile of each gene by the other. OR10G4 is expressed in the OSN Lineage almost exclusively, despite Mbnl1, the site of integration, being expressed throughout much of the olfactory epithelium.

### Figure S4

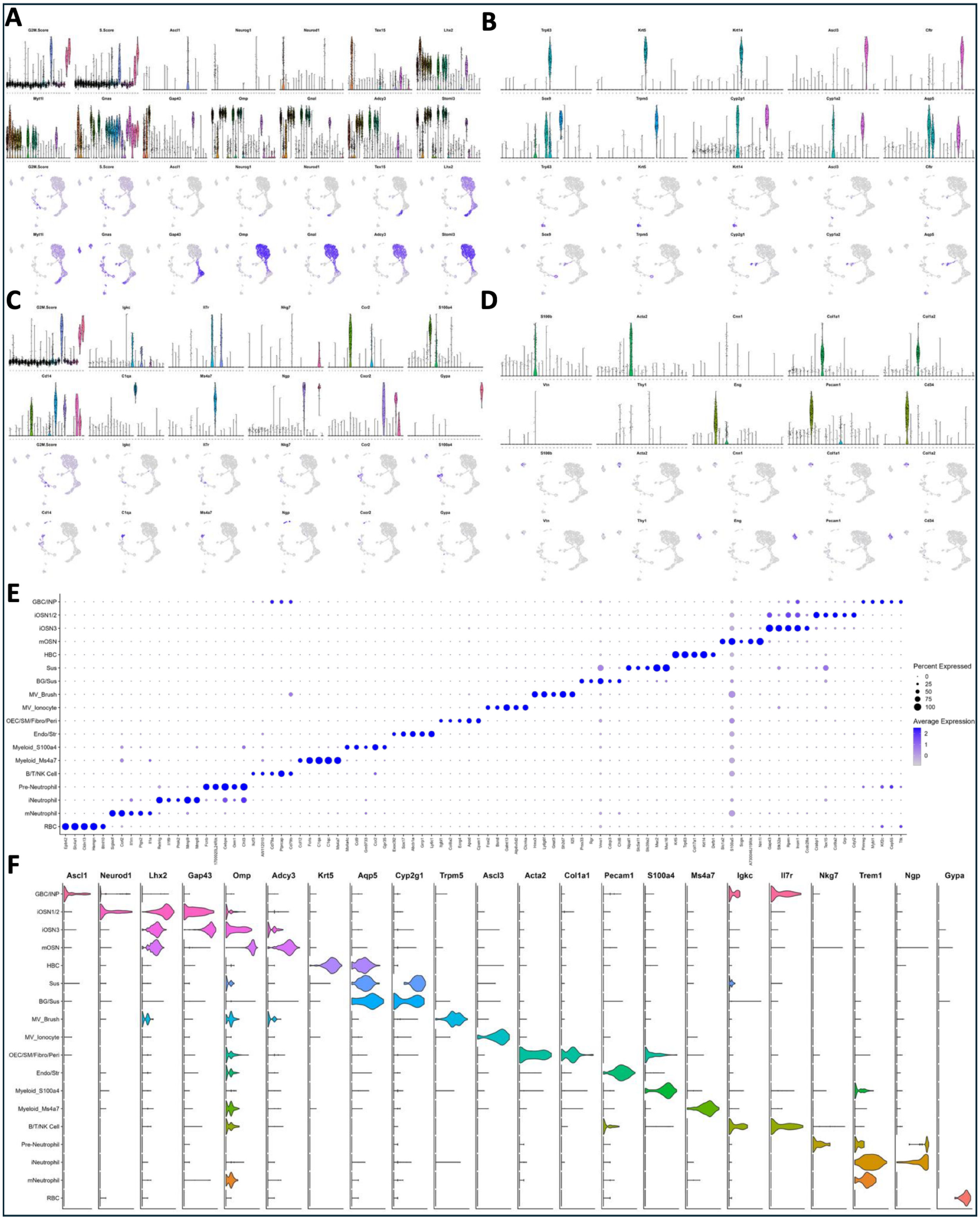
A
B
C
D
E
GBC/INP
iOSN1/2
iOSN3
mOSN
HBC
Sus
BG/Sus
MV_Brush
MV_Ionocyte
OEC/SM/Fibro/Peri
Endo/Str
Myeloid_S100a4
Myeloid_Ms4a7
B/T/NK Cell
Pre-Neutrophil
iNeutrophil
mNeutrophil
RBC
Percent Expressed
0
25
50
75
100
Average Expression
2
1
0
F
Ascl1
Neurod1
Lhx2
Gap43
Omp
Adcy3
Krt5
Aqp5
Cyp2g1
Trpm5
Ascl3
Acta2
Col1a1
Pecam1
S100a4
Ms4a7
Igkc
Il7r
Nkg7
Trem1
Ngp
Gypa

**Figure S4**, related to Figure 4. Identifying and labeling MOE clusters.

(**A-D**). Violin Plot and UMAP plot pairs show the expression profile of marker genes expressed in specific cells of the Main Olfactory Epithelium. (**A**). OSN Lineage cells: Globose Basal Cells (GBCs), Immediate Neuronal Precursors (INPs), immature OSNs (iOSNs 1-3), and Mature OSNs (mOSNs). (**B**). Epithelial cells other than OSN Lineage cells: Horizontal Basal Cells (HBCs), Sustentacular Cells (Sus), Bowman's Gland Cells (BG), and Microvillar cells (MV). (**C**). Hematopoietic and Immune cells: Various Lymphoid cells (B/T/BK Cells), Myeloid cells, Neutrophils, and Red Blood Cells (RBCs). (**D**). Lamina-based support cells: Endothelial/Stromal (Endo/Str) and Olfactory Ensheathing Cells/Smooth Muscle/Fibroblasts/Pericytes (OEC/SM/Fibro/Peri). (**E**). Dot Plot shows the top 5 differentially expressed genes in each cluster used to help identify each cluster. (**F**). Violin Plots for selected genes associated with each cell type. Labels were adequate to isolate OSN Lineage Cells.

### Figure S5

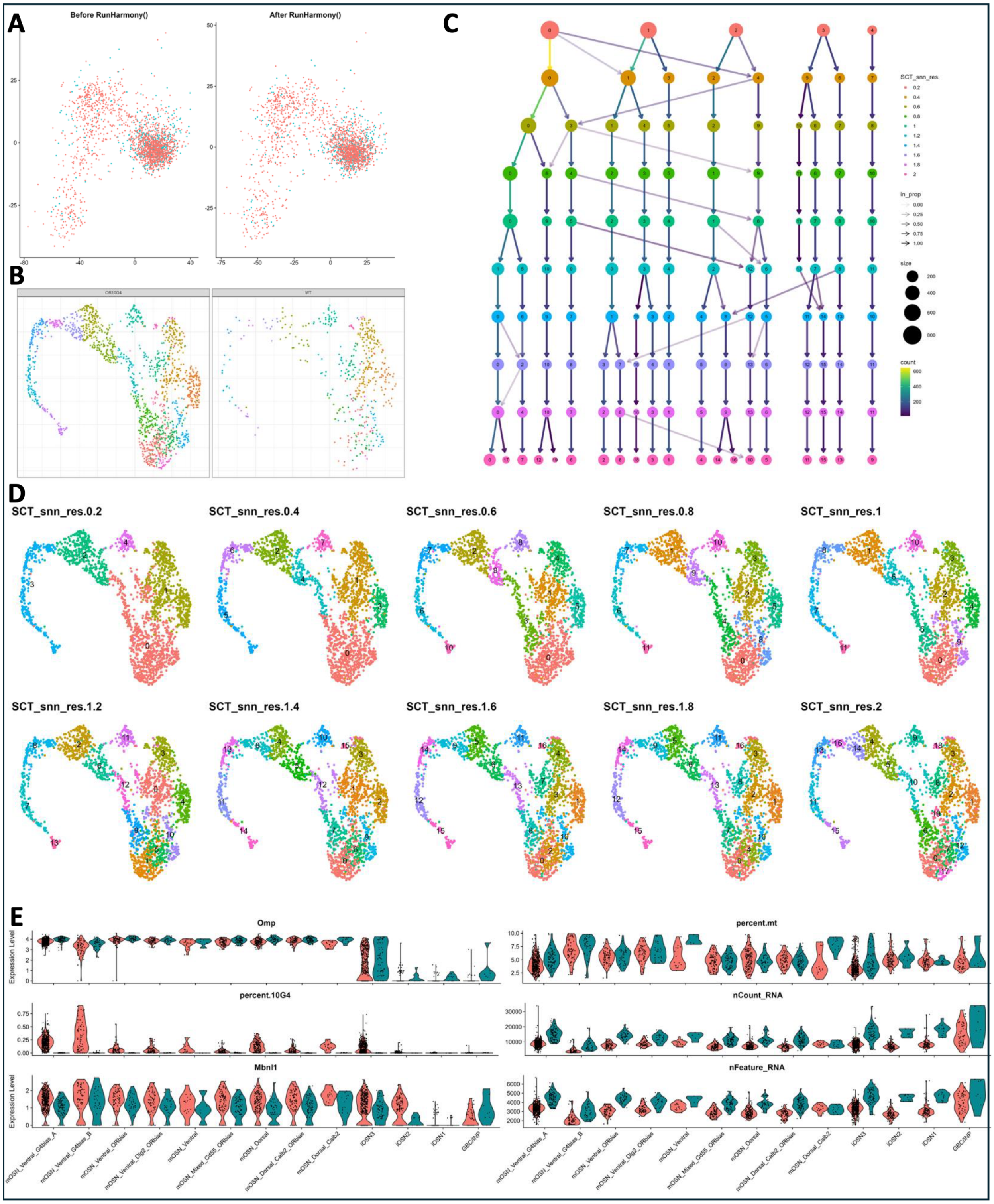
A
Before RunHarmony()
After RunHarmony()
B
OR10G4
WT
C
SCT_snn_res.
in_prop
size
count
D
SCT_snn_res.0.2
SCT_snn_res.0.4
SCT_snn_res.0.6
SCT_snn_res.0.8
SCT_snn_res.1
SCT_snn_res.1.2
SCT_snn_res.1.4
SCT_snn_res.1.6
SCT_snn_res.1.8
SCT_snn_res.2
E
Omp
percent.mt
percent.10G4
nCount_RNA
Mbnl1
nFeature_RNA
Expression Level
mOSN_Ventral_G4bias_A
mOSN_Ventral_G4bias_B
mOSN_Ventral_ORbias
mOSN_Ventral_Dlg2_ORbias
mOSN_Ventral
mOSN_Mixed_Cd55_ORbias
mOSN_Dorsal
mOSN_Dorsal_Calb2_ORbias
mOSN_Dorsal_Calb2
iOSN3
iOSN2
iOSN1
GBC/INP

**Figure S5**, related to Figure 4: Generating an OSN Lineage-only dataset.

(**A**). The Harmony package was used to combine the two datasets, showing a subtle merging of genotype-specific groups across PCA space. (**B**). UMAP shows the contribution of OR10G4 and WT samples across the various cell clusters. The two samples do not contribute equally to the various clusters. (**C**). Cluster Tree generated to evaluate cluster clarity. Highest resolution selected to better identify Genotype-specific contributions. (**D**). UMAP visualization of how resolution impacts the number of clusters and cluster splits. (**E**). Split by Genotype, violin plots for Omp expression to evaluate lineage development, OR10G4 and Mbnl1 expression to confirm the dissociation of OR10G4's expression profile from the integration locus Mbnl1's expression, and several sample properties to assess cluster-specific and genotype-specific variation.

### Figure S6

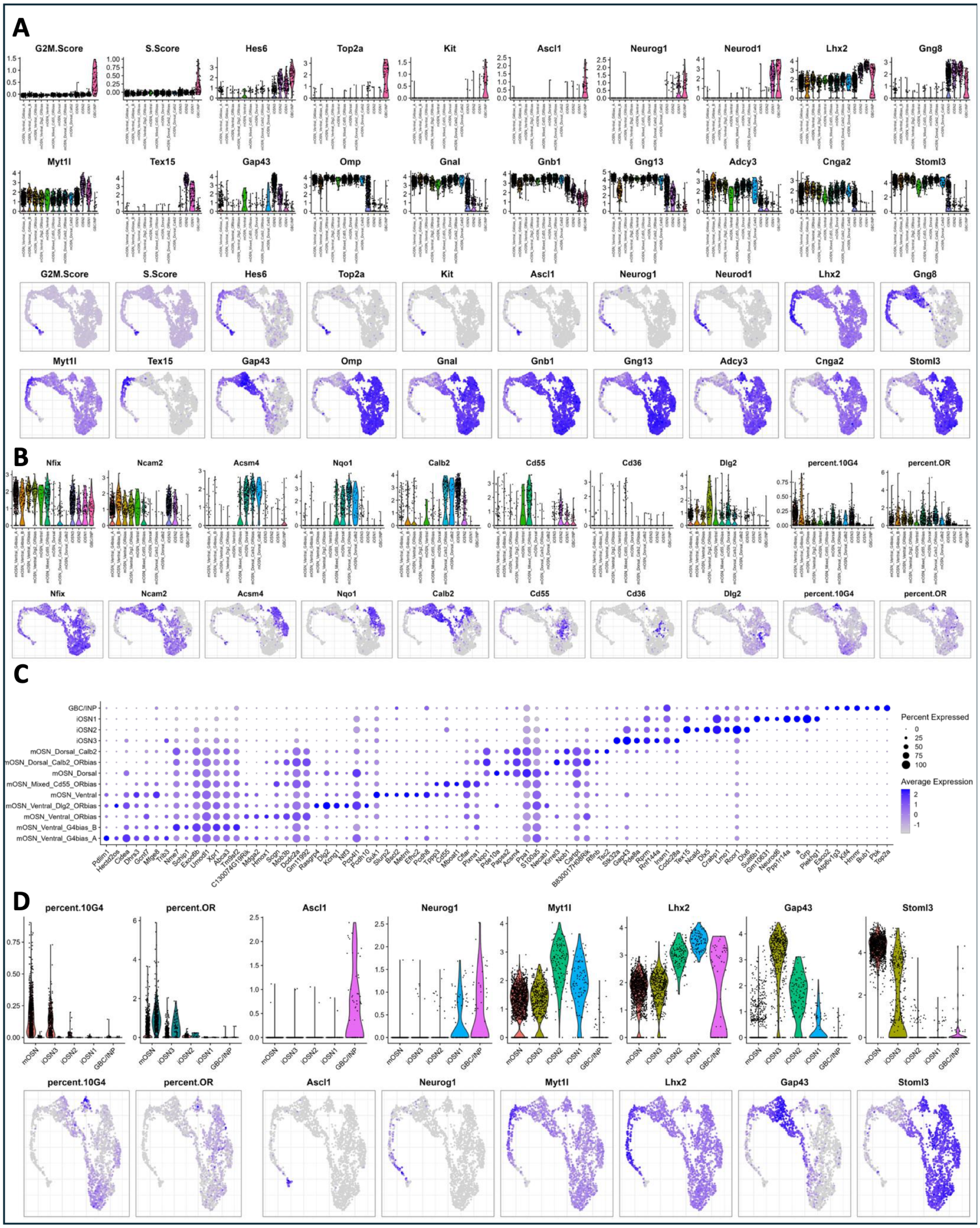
A
G2M.Score
S.Score
Hes6
Top2a
Kit
Ascl1
Neurog1
Neurod1
Lhx2
Gng8
Myt1l
Tex15
Gap43
Omp
Gnal
Gnb1
Gng13
Adcy3
Cnga2
Stoml3
B
Nfix
Ncam2
Acsm4
Nqo1
Calb2
Cd55
Cd36
Dlg2
percent.10G4
percent.OR
C
GBC/INP
iOSN1
iOSN2
iOSN3
mOSN_Dorsal_Calb2
mOSN_Dorsal_Calb2_ORbias
mOSN_Dorsal
mOSN_Mixed_Cd55_ORbias
mOSN_Ventral
mOSN_Ventral_Dlg2_ORbias
mOSN_Ventral_ORbias
mOSN_Ventral_G4bias_B
mOSN_Ventral_G4bias_A
Percent Expressed
Average Expression
D
percent.10G4
percent.OR
Ascl1
Neurog1
Myt1l
Lhx2
Gap43
Stoml3
mOSN
iOSN3
iOSN2
iOSN1
GBC/INP

**Figure S6**, related to Figure 4: Identifying specific OSN Lineage clusters and likely subdivisions of OSN Lineage cell types.

(**A**). Pairs of Violin Plots and UMAP plots that show the expression profile of genes expressed in specific cells of the OSN Lineage. (**B**). Pairs of Violin Plots and UMAP plots that show the expression profile of genes known to define subsets of OSNs. (**C**). Dot Plot shows the top 5 differentially expressed genes (DEGs) in each labeled cluster to help validate cluster labels and identify additional cluster-specific DEGs. (**D**). Using simplified cluster labels, Violin and UMAP plots pairs that show the expression profile of OR10G4 and OR genes split by Genotype (Two Left-most Plots) and the OSN Lineage markers used to identify cell types (Remaining Plots).

### Figure S7

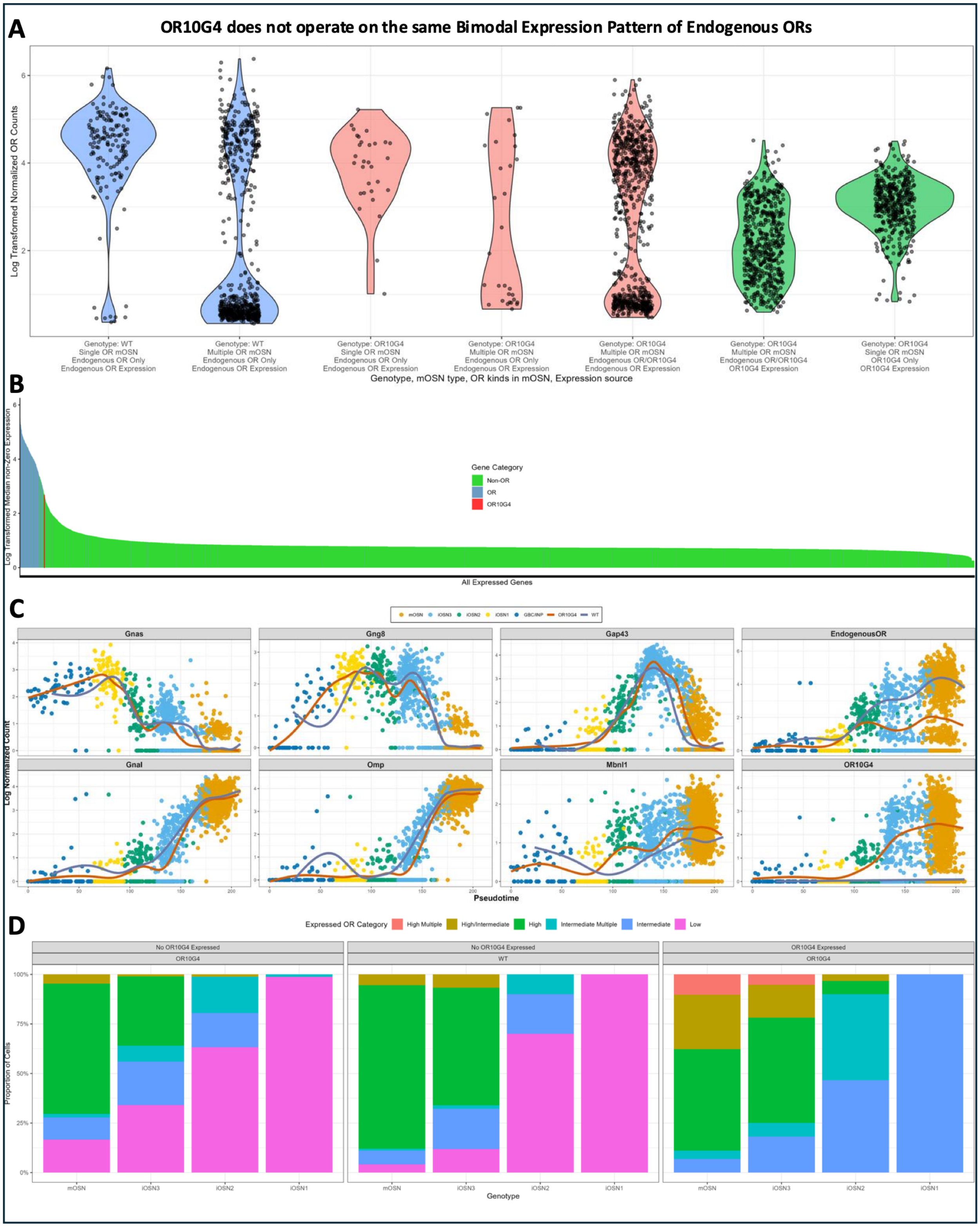
A
OR10G4 does not operate on the same Bimodal Expression Pattern of Endogenous ORs
Log Transformed Normalized OR Counts
Genotype: WT Single OR mOSN Endogenous OR Only Endogenous OR Expression
Genotype: WT Multiple OR mOSN Endogenous OR Only Endogenous OR Expression
Genotype: OR10G4 Single OR mOSN Endogenous OR Only Endogenous OR Expression
Genotype: OR10G4 Multiple OR mOSN Endogenous OR Only Endogenous OR Expression
Genotype: OR10G4 Multiple OR mOSN Endogenous OR/OR10G4 Endogenous OR Expression
Genotype: OR10G4 Multiple OR mOSN Endogenous OR/OR10G4 OR10G4 Expression
Genotype: OR10G4 Single OR mOSN OR10G4 Only OR10G4 Expression
Genotype, mOSN type, OR kinds in mOSN, Expression source
B
Log Transformed Median non-Zero Expression
Gene Category
Non-OR
OR
OR10G4
All Expressed Genes
C
mOSN
iOSN3
iOSN2
iOSN1
GBC/INP
OR10G4
WT
Gnas
Gng8
Gap43
EndogenousOR
Gnal
Omp
Mbnl1
OR10G4
Log Normalized Count
Pseudotime
D
Expressed OR Category
High Multiple
High/Intermediate
High
Intermediate Multiple
Intermediate
Low
No OR10G4 Expressed
OR10G4
No OR10G4 Expressed
WT
OR10G4 Expressed
OR10G4
Proportion of Cells
mOSN
iOSN3
iOSN2
iOSN1
Genotype

**Figure S7**, related to Figure 4. OR10G4 has a distinct count profile with lower counts that tends to violate Singular OR expression.

(**A**). A comparison between the expression of OR genes and OR10G4 in mOSNs of the WT and OR10G4 sample, further split by OR status profiles. Mature OSNs in the WT sample (blue) can have singular or multiple OR genes expressed. Endogenous OR expression in the OR10G4 sample (red) can be singular or multiple OR with or without OR10G4. OR10G4 expression is limited to the OR10G4 sample (green) and exists as either singular or alongside at least one other OR gene. **(B**). Ranking of all detected genes in mature OSNs based on the median of non-zero counts across all mOSNs. (**C**). Pseudotime evaluation of specific gene expression grouped by Genotype and colored by clustered cell type. Mbnl1 expression appears slightly earlier and elevated in the OR10G4 sample, suggesting that the integration of OR10G4 into the Mbnl1 locus elevated Mbnl1 expression during the timepoints when the OR10G4 gene is active. (**D**). Proportion bar plot comparing OR expression state per OSN between Genotypes across four developmental stages, further split by whether OR10G4 is expressed in those cells. The increase in non-singular OR expression in OR10G4 OSNs is almost entirely due to OR10G4 expression. See Figure 4E.

### Figure S8

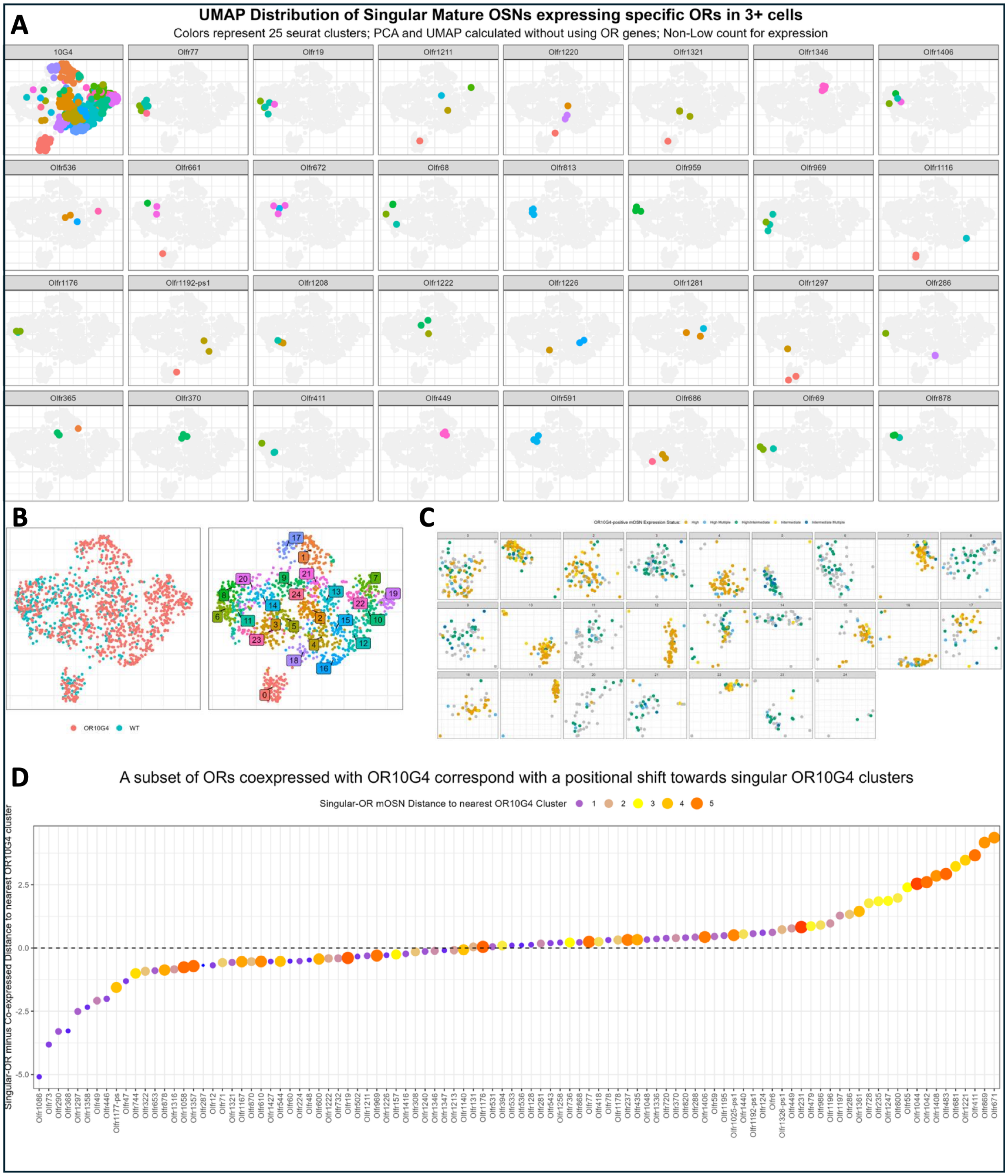
A
UMAP Distribution of Singular Mature OSNs expressing specific ORs in 3+ cells
Colors represent 25 seurat clusters; PCA and UMAP calculated without using OR genes; Non-Low count for expression
10G4
Olfr77
Olfr19
Olfr1211
Olfr1220
Olfr1321
Olfr1346
Olfr1406
Olfr536
Olfr661
Olfr672
Olfr68
Olfr813
Olfr959
Olfr969
Olfr1116
Olfr1176
Olfr1192-ps1
Olfr1208
Olfr1222
Olfr1226
Olfr1281
Olfr1297
Olfr286
Olfr365
Olfr370
Olfr411
Olfr449
Olfr591
Olfr686
Olfr69
Olfr878
B
OR10G4
WT
C
D
A subset of ORs coexpressed with OR10G4 correspond with a positional shift towards singular OR10G4 clusters
Singular-OR mOSN Distance to nearest OR10G4 Cluster 1 2 3 4 5
Singular-OR minus Co-expressed Distance to nearest OR10G4 cluster

**Figure S8**, related to Figure 4. OR-specific mOSN “mini-clusters” can be shifted towards OR10G4 clusters by OR10G4 co-expression.

(**A**). As established for OSN Lineage cells, mature OSNs were subset and clustered sans OR genes to generate an OR transcript-agnostic UMAP. ORs expressed in singularity in at least 3 mOSNs were identified and marked on separate UMAPs to evaluate if same-OR mOSNs cluster together in UMAP space. (**B**). UMAP plots showing (Left) OR10G4 and WT cells unevenly distributed through the space and (Right) the numerical labels for individual clusters. (**C**). UMAP evaluation of categorized OR10G4 expression split across the 25 clusters generated during the clustering process. Singular OR10G4 gene expression is limited to a subset of clusters, as seen in (A). Grey circles do not meet the threshold for OR10G4 expression. Clusters with substantial singular OR10G4 mOSNs (>=10%) are defined as “Singular OR10G4 clusters”. (**D**). The mean UMAP position of Singular OR and OR/OR10G4 “mini-clusters”, and Singular OR10G4 clusters was determined and used to evaluate OR-specific changes in the distance between their associated mini-clusters and the nearest OR10G4 cluster. A point near the dashed line at Zero means the OR/OR10G4 mini-cluster did not appear any closer or further away from any OR10G4 cluster than from the “baseline” Singular-OR mini-cluster. Positive and Negative values represent positional shifts closer and further away, respectively, from OR10G4 clusters in the context of OR10G4 co-expression. Size and color are linked to better emphasize Singular-OR proximity to the nearest OR10G4 cluster.

### Figure S9

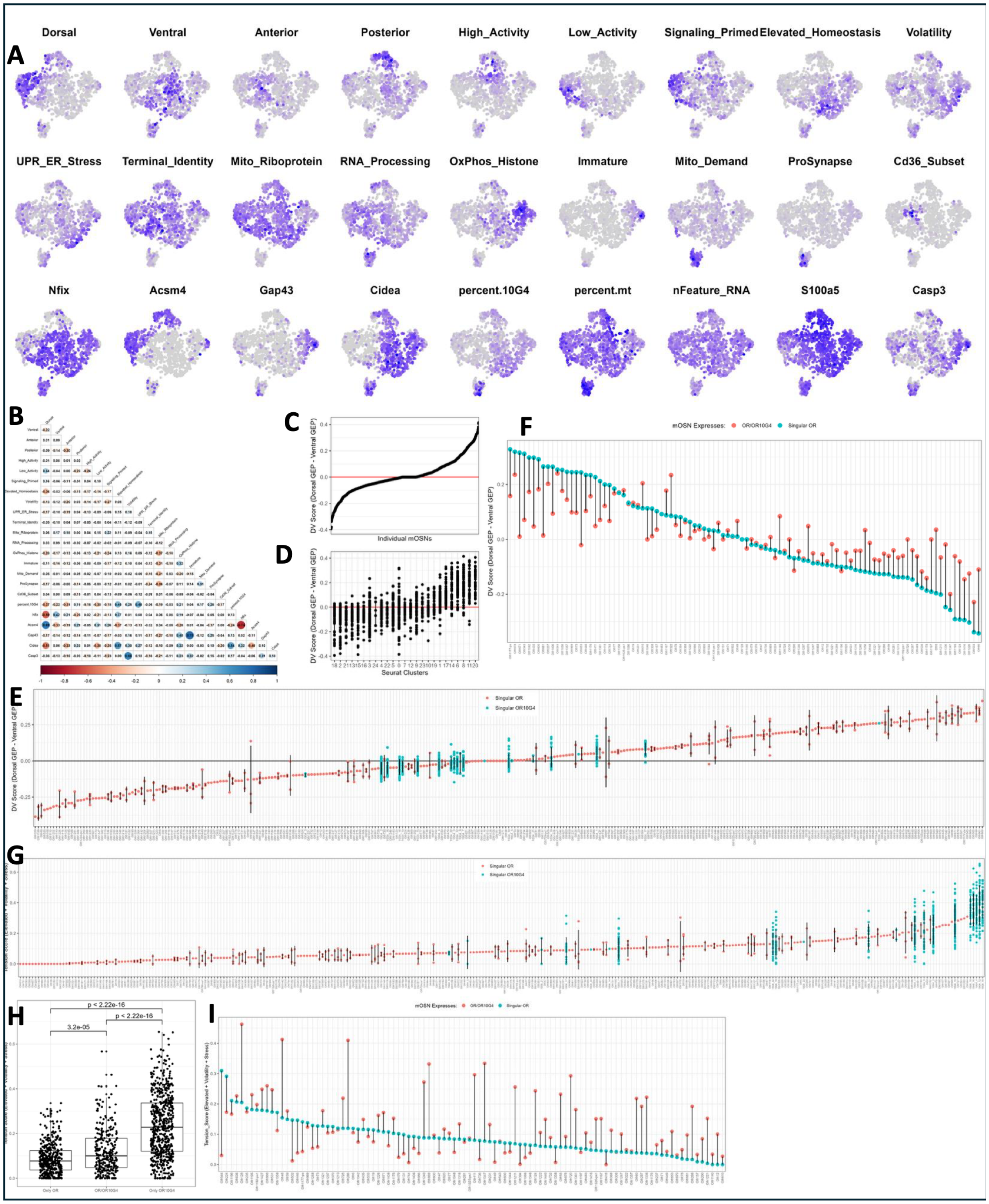
A
Dorsal
Ventral
Anterior
Posterior
High_Activity
Low_Activity
Signaling_Primed
Elevated_Homeostasis
Volatility
UPR_ER_Stress
Terminal_Identity
Mito_Riboprotein
RNA_Processing
OxPhos_Histone
Immature
Mito_Demand
ProSynapse
Cd36_Subset
Nfix
Acsm4
Gap43
Cidea
percent.10G4
percent.mt
nFeature_RNA
S100a5
Casp3
B
C
DV Score (Dorsal GEP - Ventral GEP)
Individual mOSNs
D
Seurat Clusters
F
mOSN Expresses: OR/OR10G4 Singular OR
E
Singular OR
Singular OR10G4
G
H
p < 2.22e-16
3.2e-05
p < 2.22e-16
I

**Figure S9**, related to Figure 4. Gene Expression Program analysis, as with the UMAP distance metric, shows that OR10G4 co-expression can cause a shift towards an OR10G4-like GEP profile.

(**A**). Mature OSN gene counts were processed via consensus non-negative matrix factorization (cNMF) to produce 24 gene expression programs (GEPs). GEPs 19-24 were active in too few cells and are not shown. UMAP plots display GEP usage across mOSNs. GEPs were evaluated and then labeled based on expression of marker genes like Nfix and Acsm4, Gene Ontology Enrichment Analysis and AI summaries of top loading score genes and compared with published OSN GEPs. (**B**). Correlation heatmap for GEPs and selected genes. (**C**). The DV Score is calculated as the difference between Dorsal and Ventral GEP usage, as in Tsukahara et al., 2021. The DV Score for all individual cells is arranged from smallest (most Ventral) to largest (most Dorsal). Red line is a visual aid. (**D**). The average DV Score for each mOSN cluster was used to arrange clusters and the DV scores for all constituent cells grouped by cluster were plotted. The lowest DV Score clusters have nearly zero overlap with the highest DV Score clusters. (**E**). The average DV Score for Singular OR mOSNs grouped by expressed OR, including OR10G4 suffixed by cluster (See S8C), shows a continuous spread of values, though ORs with greater than one mOSN show a clear overlap in said scores with many other OR-associated DV scores. OR10G4 DV Scores sit between approximately -0.125 and 0.125, crowding around or slightly negative of zero. (**F**). The same 100 OR groups from Supplemental Figure 8D were evaluated. The DV Score for mOSNs that express an OR alongside OR10G4 tends to shift from their Singular-OR DV score towards zero where Singular OR10G4 DV scores appear. (**G**). A "Tension" Score was calculated for each cell based on the sum of the Elevated_Homeostasis, Volatility, and UPR_ER_Stress GEPs, three GEPs that correlate positively with OR10G4 expression. Most Singular OR mOSNs have a low Tension Score, while most, but not all Singular OR10G4 mOSN groups have a high Tension Score. (**H**). The Tension Score for Singular OR, Singular OR10G4, and OR/OR10G4 mOSNs are all significantly different. Corrections for multiple comparisons not applied. (**I**). As in (F), Tension Scores were compared for ORs that have Singular OR and OR/OR10G4 mOSNs. Over half of the ORs were associated with an increase in their score when co-expressing OR10G4, though the effect is most prevalent for ORs that start with a low Tension Score in singular-OR mOSNs.

### Figure S10

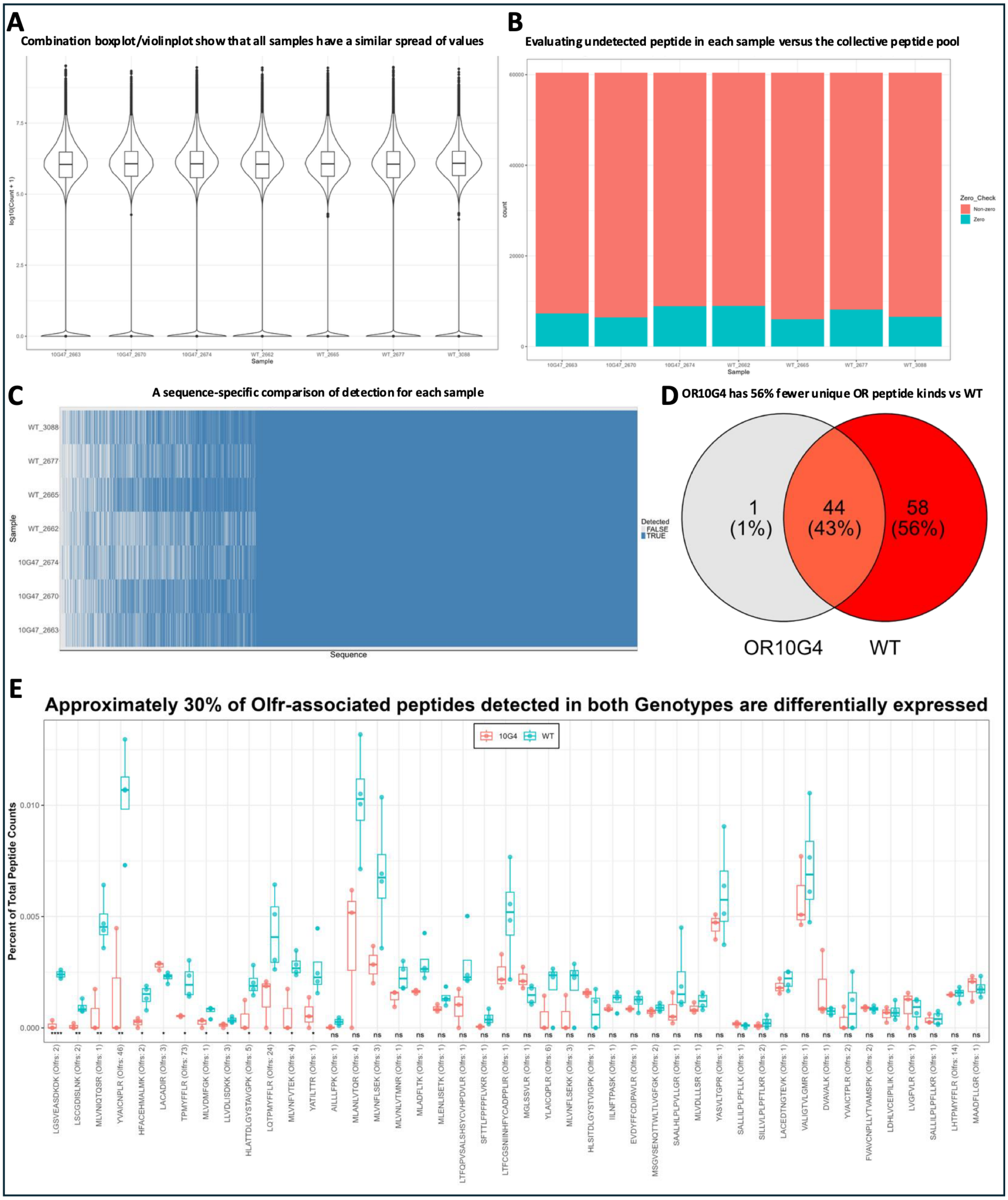
A
Combination boxplot/violinplot show that all samples have a similar spread of values
log10(Count + 1)
Sample
10G47_2663
10G47_2670
10G47_2674
WT_2662
WT_2665
WT_2677
WT_3088
B
Evaluating undetected peptide in each sample versus the collective peptide pool
count
Zero_Check
Non-zero
Zero
C
A sequence-specific comparison of detection for each sample
Sample
Sequence
Detected
FALSE
TRUE
D
OR10G4 has 56% fewer unique OR peptide kinds vs WT
1 (1%)
44 (43%)
58 (56%)
OR10G4
WT
E
Approximately 30% of Olfr-associated peptides detected in both Genotypes are differentially expressed
10G4
WT
Percent of Total Peptide Counts
LGSVEASDKDK (Olfrs: 2)
LSCGDISLNK (Olfrs: 2)
MLVNIQTQSR (Olfrs: 1)
YVAICNPLR (Olfrs: 46)
HFACEHMALMK (Olfrs: 2)
LACADIR (Olfrs: 3)
TPMYFFLR (Olfrs: 73)
MLVDMFGK (Olfrs: 1)
LLVDLISDKK (Olfrs: 3)
HLATTDLGYSTAVGPK (Olfrs: 5)
LQTPMYFFLR (Olfrs: 24)
MLVNFVTEK (Olfrs: 4)
YATILTTR (Olfrs: 1)
AILLLFPK (Olfrs: 1)
MLANLVTQR (Olfrs: 4)
MLVNFLSEK (Olfrs: 3)
MLVNLVTMNR (Olfrs: 1)
MLADFLTK (Olfrs: 1)
MLENLISETK (Olfrs: 1)
LTFQPVSALSHSYCVHPDVLR (Olfrs: 1)
SFTTLFPFPFLVKR (Olfrs: 1)
LTFCGSNIINHFYCADPPLIR (Olfrs: 1)
MGLSSVLR (Olfrs: 1)
YLAICQPLR (Olfrs: 6)
MLVNFLSEKK (Olfrs: 3)
HLSITDLGYSTVIGPK (Olfrs: 1)
IILNFTPASK (Olfrs: 1)
EVDYFFCDIPAVLR (Olfrs: 1)
MSGVSENQTTWLTLVGFGK (Olfrs: 2)
SAALHLPLPVLLGR (Olfrs: 1)
MLVDLLLSR (Olfrs: 1)
YASVLTGPR (Olfrs: 1)
SALLILPLPFLLK (Olfrs: 1)
SILLVLPLPFTLKR (Olfrs: 2)
LACEDTNGTEVK (Olfrs: 1)
VALIGTVLGMR (Olfrs: 1)
DVAVALK (Olfrs: 1)
YVAICTPLR (Olfrs: 2)
FVAVCNPLLYTVAMSPK (Olfrs: 2)
LDHLVCEIPILIK (Olfrs: 1)
LVGFVLR (Olfrs: 1)
SALLILPLPFLLKR (Olfrs: 1)
LHTPMYFFLR (Olfrs: 14)
MAADFLLGR (Olfrs: 1)

**Figure S10**, related to Figure 5. OR10G4 and WT samples share similar quality control values but differ in detection of OR peptides.

(**A**). Violin Plot / Box Plot combination shows a similar distribution of count values for all samples. Sample 2677 is significantly different from 2670 and 3088 after multiple corrections (not shown), likely due to technical and biological variance. (**B**). Approximately 60,000 unique peptide sequences were detected between all samples, though each individual sample was missing approximately 7500 peptides from the collective sequencing result. (**C**). A visualization of ubiquitous and undetected peptides for each sample. (**D**). As an alternative to the default peptide-to-protein aggregate value, the number of unique OR peptides detected in each genotype was compared using a Venn Diagram. 103 OR protein-associated peptides are present in the mass spec sequencing result with only a single peptide unique to OR10G4. (**E**). Out of 44 OR peptides detected in both genotypes, 13 are differentially expressed with several others showing a similar trend. T test results were not corrected for multiple comparisons. Counts normalized to individual sample totals. Significance: *p <= 0.05, **p <= 0.01, ***p <= 0.001, ****p <= 0.0001, ns = Not Significant

### Figure S11

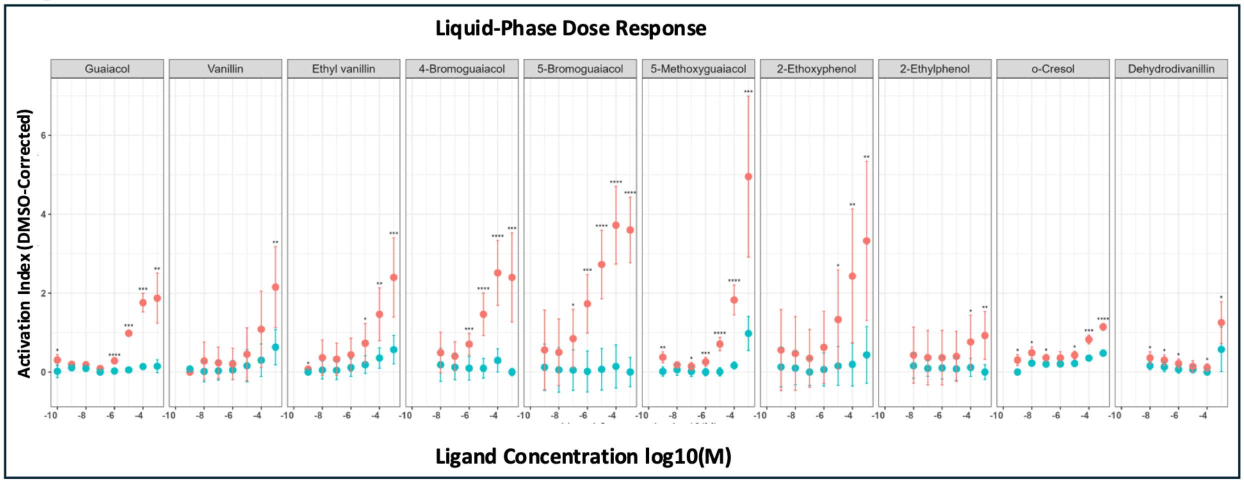
Liquid-Phase Dose Response
Guaiacol
Vanillin
Ethyl vanillin
4-Bromoguaiacol
5-Bromoguaiacol
5-Methoxyguaiacol
2-Ethoxyphenol
2-Ethylphenol
o-Cresol
Dehydrodivanillin
Activation Index (DMSO-Corrected)
Ligand Concentration log10(M)

**Figure S11**, related to Figure 5. Several odor molecules perform as well or better than guaiacol in the cilia cAMP assay. Ligand Dose Responses are induced by various odors in OR10G4 vs WT-derived cilia. Each ligand was tested 3 times. No p value corrections applied. Error bars: Activation Index mean +/- sd. Significance: *p <= 0.05, **p <= 0.01, ***p <= 0.001, ****p <= 0.0001, ns = Not Significant

**Supplemental Tables**

**Supplemental Table S1**, related to Figure 2. Comparing Bulk RNA-seq results to RT-qPCR analysis limited to the small OR subset tested with RT-qPCR. The RNA-seq results are nearly identical to the Line 7 RT-qPCR. The same RNA samples were used in both procedures; thus, the results are not unexpected but validate the use of small-throughput tests.
**Supplemental Table S2**, related to Figure 2. Directly from Tsukahara et al. 2021, who defined the Dorsal/Ventral Index values for each OR when possible.
**Supplemental Table S3,** related to Figure 4. Table that categorizes the expression of each OR gene, including OR10G4, based on Genotype, Singular-OR vs. multiple-OR OSN, presence of OR10G4 expression, expression category for each count instance, and how those count instances are distributed throughout the various categories of OSN-OR kinds. Most Low counts are absorbed into OSNs with non-Low counts as background or basal expression, though Low-only OSNs exist and are named as such.
**Supplemental Table S4**, Related to Figure 4. GEP tables.
**Supplemental Table S5**, Related to Figure 5. A summary of each odor molecule used in this manuscript.
**Supplemental Table S6**, Related to Figure 1. RT-qPCR experimental design details.
**Supplemental Table S7**, Related to Figure 1. Genotyping Primers.
