## Supplementary figures and images for "Functional Profiling of a Human Odorant Receptor Using Olfactory Cilia Links its Activation to Odor Quality"

### Supplemental Table 1

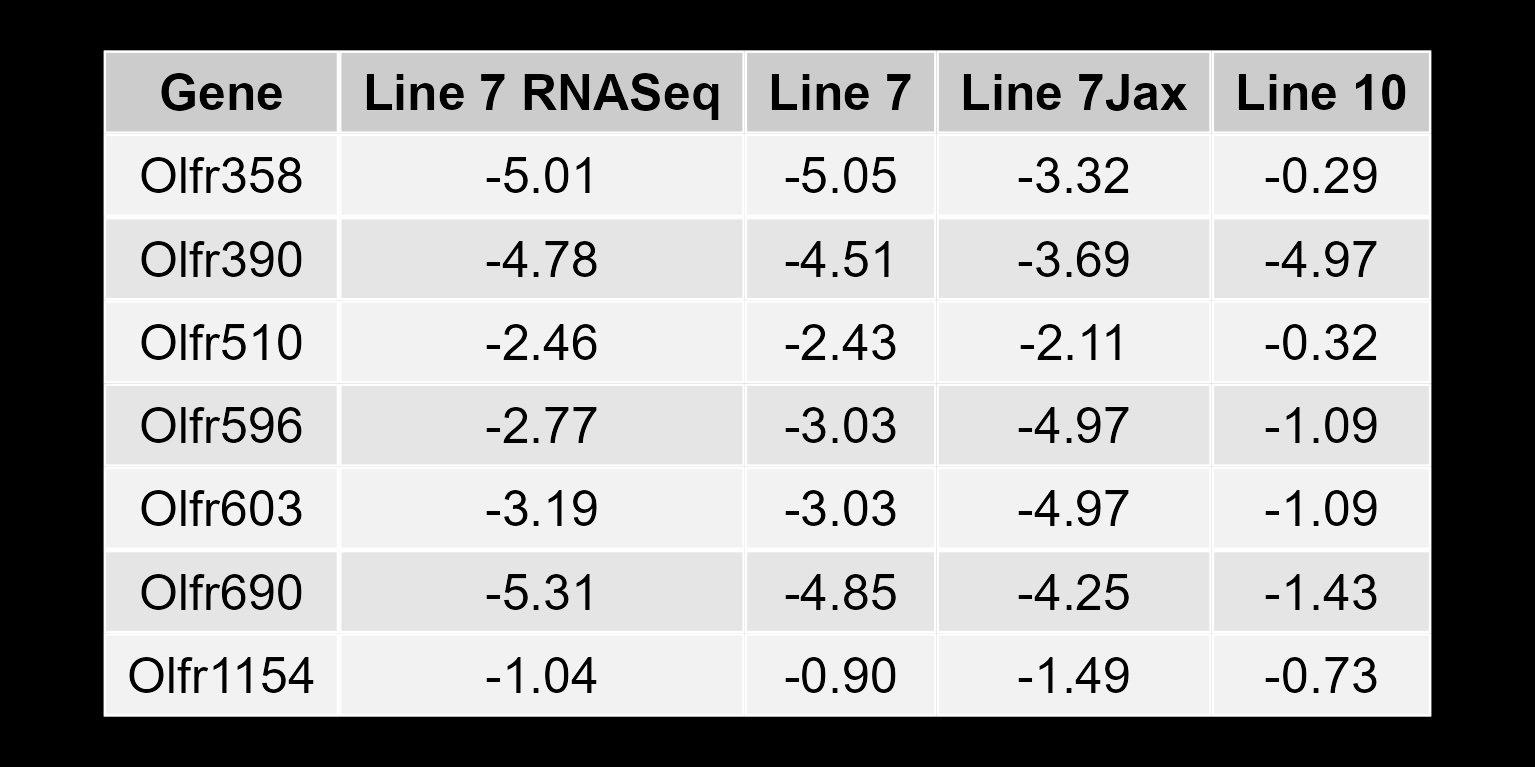
