## Supplemental Table 5 for "Functional Profiling of a Human Odorant Receptor Using Olfactory Cilia Links its Activation to Odor Quality"

|  | **Trivial Name** | **IUPAC name**  **(non-standard arrangements)** | **Structure** | **Mean A.I. (minus WT)** | **Median A.I. (minus WT)** | **OR10G4 EC50, Liquid** | **Vapor Pressure (Mackay Method) Epi Suite** | **Odor Threshold, Median (ng/Lair)** | **Odor qualities, Aggregate** | **References: Odor Threshold and qualities** |
| --- | --- | --- | --- | --- | --- | --- | --- | --- | --- | --- |
| 1 | Guaiacol | 2-Methoxyphenol | 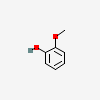 | 1.1888 | 1.1378 | 0.00001109 | 1.10 | 0.084 | smoky, vanilla, ham | Schranz, et al. 2017. |
| 2 | Vanillin | 4-Formyl-2-methoxyphenol | 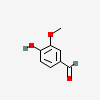 | 0.2729 | 0.1915 | 0.0003240 | 0.00871 | 8 | sweet, vanilla, creamy, phenolic, chocolate | de-la-Fuente-Blanco & Ferreira. 2020;  https://www.thegoodscentscompany.com/data/rw1011712.html |
| 3 | 4-Chloroguaiacol | 4-Chloro-2-methoxyphenol | 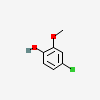 | 0.9855 | 0.5362 | NA | 0.189 | 0.35 | *sweet*, vanilla | Juhlke, et al. 2017 |
| 4 | 4-Bromoguaiacol | 4-Bromo-2-methoxyphenol | 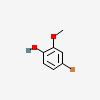 | 2.5704 | 1.8628 | 0.000009813 | 0.0470 | 0.029 | *vanilla, sweet*, smoky | Juhlke, et al. 2017 |
| 5 | 5-Bromoguaiacol | 5-Bromo-2-methoxyphenol | 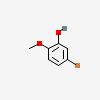 | 3.4474 | 2.7897 | 0.000002425 | 0.0470 | 0.0023 | *smoky*, sweet | Juhlke, et al. 2017 |
| 6 | 4-Iodoguaiacol | 4-Iodo-2-methoxyphenol | 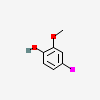 | 0.4416 | 0.2014 | NA | 0.00946 | 4.1 | *vanilla*, smoky, sweet | Juhlke, et al. 2017 |
| 7 | 4-Fluoroguaiacol | 4-Fluoro-2-methoxyphenol | 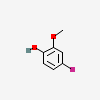 | 1.2417 | 0.8097 | NA | 1.10 | NA | NA |  |
| 8 | 5-Fluoroguaiacol | 5-Fluoro-2-methoxyphenol | 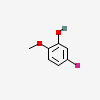 | 2.5907 | 2.0552 | NA | 1.10 | NA | NA |  |
| 9 | 4-Methylguaiacol | 4-Methyl-2-methoxyphenol | 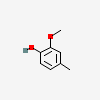 | 1.1225 | 0.6508 | NA | 0.476 | 1.4 | vanilla, sweet, ham, smoky | Schranz, et al. 2017. |
| 10 | 4-Methoxyguaiacol | 2,4-Dimethoxyphenol | 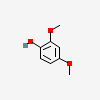 | 0.8374 | 0.4299 | NA | 0.0868 | 1.0 | clove, sweet, smoky, vanilla, ham | Schranz, et al. 2017. |
| 11 | 5-Methoxyguaiacol | 2,5-Dimethoxyphenol | 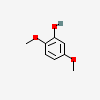 | 2.7704 | 2.4289 | 0.003572* | 0.0868 | 0.00018 | sweet, clove, vanilla | Schranz, et al. 2017. |
| 12 | Dehydrodivanillin | 2,2'-Dimethoxy-4,4'-Diformyl-6,6'-Biphenol | 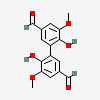 | 0.2024 | 0.1280 | NA** | 0.0000000188 | NA | fruity, vanilla | https://www.thegoodscentscompany.com/data/rw1407621.html |
| 13 | Acetovanillone | 4-Acetyl-2-methoxyphenol | 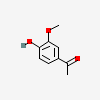 | 0.1125 | 0.0583 | NA | 0.00232 | NA | sweet, vanilla | https://www.thegoodscentscompany.com/data/rw1057511.html |
| 14 | Vanillyl butyl ether | 4-(Butoxymethyl)-  2-methoxyphenol | 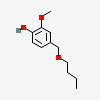 | 0.0780 | -0.0119 | NA | 0.00173 | NA | vanilla, fruity | https://www.thegoodscentscompany.com/data/rw1038461.html |
| 15 | Eugenol | 4-Allyl-2-methoxyphenol | 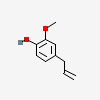 | 0.2457 | 0.0728 | NA | 0.0976 | NA | sweet, spicy, clove, woody, phenolic | https://www.thegoodscentscompany.com/data/rw1004991.html |
| 16 | 2-Hydroxy-4-Methoxybenzaldehyde | 2-Formyl-5-methoxyphenol | 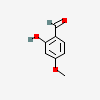 | 0.2109 | 0.0780 | NA | 0.0182 | NA | vanilla | https://www.thegoodscentscompany.com/data/rw1027851.html |
| 17 | Ethyl Vanillin | 4-Formyl-2-ethoxyphenol | 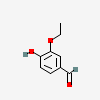 | 0.6067 | 0.4693 | 0.0003284 | 0.00613 | NA | sweet, creamy, vanilla, caramel | https://www.thegoodscentscompany.com/data/rw1002652.html |
| 18 | 2-Ethoxyphenol | 2-Ethoxyphenol | 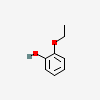 | 2.2913 | 1.6161 | 0.00004661 | 0.681 | NA | phenolic, medicinal | Lo, et al. 2008 |
| 19 | o-Cresol | 2-Methylphenol | 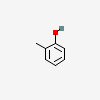 | 0.3837 | 0.1810 | 0.00008251 | 2.10 | NA | musty, phenolic, plastic, medicinal, herbal | https://www.thegoodscentscompany.com/data/rw1007721.html |
| 20 | 2-Ethylphenol | 2-Ethylphenol | 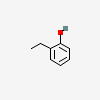 | 0.5033 | 0.2446 | 0.00006701 | 1.01 | NA | phenolic | https://www.thegoodscentscompany.com/data/rw1175631.html |
| 21 | Salicylaldehyde | 2-Formylphenol | 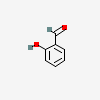 | 0.2669 | 0.0495 | NA | 1.41 | NA | medicinal, spicy, cinnamon, wintergreen, cooling | https://www.thegoodscentscompany.com/data/rw1028641.html |
| 22 | Menthoxypropanediol | 3-[[5-methyl-2-(1-methylethyl)cyclohexyl]oxy]-1,2-propanediol | 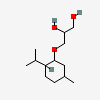 | 0.2701 | 0.0507 | NA | 0.000899 | NA | minty, phenolic, fruity, jammy, berry | https://www.thegoodscentscompany.com/data/rw1038341.html |
